# Metabolization of microbial postbiotic pentanoate drives anti-cancer CAR T cells

**DOI:** 10.1101/2024.08.19.608538

**Authors:** Sarah Staudt, Fabian Nikolka, Markus Perl, Julia Franz, Noémi Leblay, Xiaoli-Kat Yuan, M Larrayoz, Teresa Lozano, Linda Warmuth, Matthias A. Fante, Aistė Skorupskaitė, Teng Fei, Maria Bromberg, Patxi San Martin-Uriz, Juan Roberto Rodriguez-Madoz, Kai Ziegler-Martin, Nazdar Adil Gholam, Pascal Benz, Phuc-Huu Tran, Fabian Freitag, Zeno Riester, Christoph Stein-Thoeringer, Michael Schmitt, Karin Kleigrewe, Justus Weber, Kira Mangold, Patrick Ho, Hermann Einsele, Felipe Prosper, Wilfried Ellmeier, Dirk Busch, Alexander Visekruna, John Slingerland, Roni Shouval, Karsten Hiller, Juan José Lasarte, José Ángel Martinez-Climent, Patrick Pausch, Paola Neri, Marcel van den Brink, Hendrik Poeck, Michael Hudecek, Maik Luu

## Abstract

The microbiome is a complex host factor and key determinant of the outcome of antibody-based and cellular immunotherapy. Its postbiotics are a blend of soluble commensal byproducts that are released into the host environment and have been associated with the regulation of immune homeostasis, particularly through impacts on epigenetics and cell signaling.

In this study, we show that the postbiotic pentanoate is metabolized to citrate within the TCA cycle via both the acetyl- and succinyl-CoA entry points, a feature uniquely enabled by the chemical structure of the C5 aliphatic chain. We identified ATP-citrate lyase as the crucial factor that redirects pentanoate-derived citrate from the succinyl-CoA route to the nucleus, thereby linking metabolic output and histone acetylation. This epigenetic-metabolic crosstalk mitigated T cell exhaustion and promoted naive-like differentiation in pentanoate-programmed chimeric antigen receptor (CAR) T cells. The predictive and therapeutic potential of pentanoate was corroborated in two independent patient cohorts and three syngeneic models of CAR T adoptive therapy.

Our data demonstrate that postbiotics are integrated into mitochondrial metabolism and subsequently incorporated as epigenetic imprints. This bridge between microbial and mammalian interspecies communication can ultimately impact T cell differentiation and efficacy.

## Introduction

The intestinal microbiome plays a key role in determining immunotherapy outcomes across a multitude of pathophysiological conditions, including cancers. [1–3]. For example, the composition of gut microbiota has been shown to influence immune-cell activation and differentiation within tumor microenvironments (TMEs) [4, 5]. One mode of regulation involves translocation of commensal bacteria from the intestine to distal tumor sites, a finding that has inspired the design of commensal consortia that promote T cell responses during immune checkpoint inhibition (ICI) [6, 7]. Unfortunately, the complexity of interactions between individual microbial species presents a major challenge to strategies that aim to reshape the microbiome, either at the bulk or strain-specific level [8, 9]. As an alternative to defining microbiome composition, microbial metabolites (postbiotics) can be generated and delivered as effector molecules. However, the postbiotics profile of various bacterial strains can cause overlapping or neutralizing effects, demanding in-depth characterization of individual postbiotics [10, 1, 11–14].

In previous work, we have investigated the role of different short-chain fatty acids (SCFAs; a major group of commensal molecules) in inflammation, autoimmunity, and cancer [10, 15–17].

Interestingly, the same metabolite can drive opposing immunosuppressive and stimulatory effects, highlighting the potential for context- and cell type-specific modulation. One such compound is pentanoate, a rare SCFA that can be produced by lowly abundant strains such as *Megasphaera massiliensis*. While pentanoate exposure induced a regulatory phenotype in B cells to curb autoimmunity, pentanoate treatment was able to augment the anti-tumor efficacy of TCR-transgenic CD8^+^ T cells [18]. These promising results prompted us to investigate its relevance for CAR T cell patients.

In this study, we uncover a strong correlation between the abundance of pentanoate and progression-free survival (PFS) of cancer patients receiving CAR T cells. By employing metabolic tracing of ^13^C-labeled pentanoate, we revealed that the C5 molecule becomes metabolized in the tricarboxylic acid (TCA) cycle via two different entry points, contributing to citrate translocation into the nucleus and consequently histone acetylation. We used a spectrum of pre-clinical models and two independent clinical cohorts to corroborate our findings.

The data show that commensal postbiotics can drive epigenetic imprinting in T cells and confer features that cannot be replicated by currently available drugs, thereby underlining the importance of communication between commensals and immune cells for immunotherapies.

## Results

### Pentanoate is a predictor of patient survival

We evaluated the relationship between fecal SCFA levels and PFS of patients receiving CAR- T cells in a German cohort of lymphoma and myeloma patients (**Suppl. Table 1)**. Patients with high levels of pentanoate exhibited significantly better PFS compared to those with low levels of pentanoate (Hazard ratio (HR) 6.9, estimated 1-year PFS 90% vs. 41.5%; median PFS not reached vs. 310 days, **Fig. 1a**). These patients also exhibited a trend towards improved overall survival (HR 3.0, estimated 1 year overall survival (OS) 80% vs. 71.7%, median OS not reached vs. 749 days, **Fig. 1a**). Notably, other SCFAs such as butyrate, propionate and acetate could not predict PFS and OS to a similar degree (**Fig. 1c-d**). In a second cohort of US patients (n=60) with non-Hodgkin lymphoma treated with CD19- directed CAR T cells (**Suppl. Table 2),** the pre-CAR T infusion fecal pentanoate concentrations were categorized by terciles. The higher terciles had numerically higher 2-year PFS (p=0.4) and 2-year OS (p=0.4, **Fig. 1e,f**).

**Figure 1:**
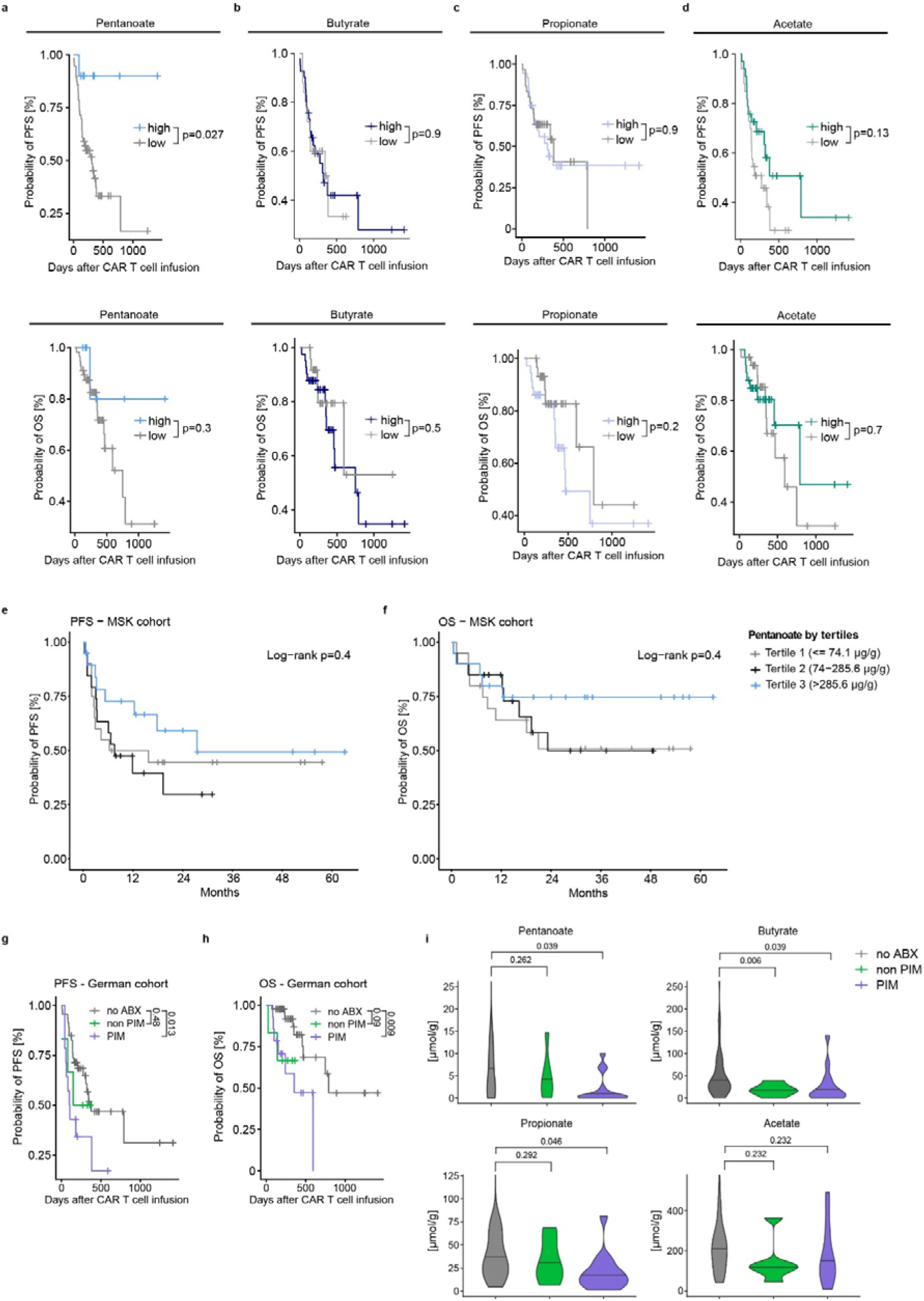
Pentanoate abundance is a predictor of clinical CAR T cell response. **a**-**d,** Kaplan-Meier curves showing PFS (upper panel) and OS (lower panel) depending on the abundance of pentanoate (**a**), butyrate (**b**), propionate (**c**) and acetate (**d**). **e** and **f**, Probability of PFS (**e**) and overall survival (**f**) in a cohort of patients receiving CAR T cell therapy depending on the pentanoate concentration. **g, h,** Probability of PFS (**g**) and overall survival (**h**) in a cohort of patients receiving CAR T cells depending on different antibiotic-(ABX)- treatment regimen. PIM (piperacillin/tazobactam, imipenem, meropenem); PFS, progression-free survival; OS, overall survival. **i,** Levels of multiple short-chain fatty acids following antibiotic exposure.

Thus, pentanoate abundance constitutes a predictive marker of CAR T cell response. Previous reports have highlighted the impact of broad-spectrum antibiotics on anti-CD19 CAR T cell therapy outcomes [8, 9]. Patients exposed to PIM group antibiotics (i.e., piperacillin/tazobactam, imipenem, meropenem) prior to CAR T cell therapy exhibited worse PFS (HR 2.66, **Fig. 1g**) and OS (HR 4.11, **Fig. 1h**) compared to patients not exposed to antibiotics. Conversely, patients receiving non-PIM antibiotics showed similar PFS (HR 1.55, **Fig. 1g**) and only a trend towards adverse OS (HR 3.95, **Fig. 1h**) compared to those not treated with antibiotics.

Examination of the association between SCFA levels and antibiotic exposure revealed that pentanoate levels were significantly reduced only in patients with PIM exposure (**Fig. 1i**). No significant reduction was observed with non-PIM antibiotics. A similar effect could be observed for the odd-chain family member propionate, albeit to a far lesser extent. However, levels of the even-chain fatty acid butyrate, were already decreased with non-PIM antibiotics. Acetate showed no significant reduction with either PIM or non-PIM antibiotics. These correlations suggest that PIM exposure substantially affects pentanoate-producing commensals and thereby PFS.

### Pentanoate modulates the CAR T cell phenotype

We hypothesized that the superior PFS in lymphoma and multiple myeloma (MM) might be due to pentanoate-mediated immunostimulation, potentially resulting in a less hostile host environment and greater CAR T cell function. To investigate whether pentanoate could enhance the anti-tumor efficacy of CAR T cells in a clinically relevant setting, we incorporated pentanoate treatment into the production process. A potential advantage of pentanoate might be its ability to acce,erate epigenetic reprogamming. Thus, we designed a shortened production process to benefit from early chromatin remodeling and to better retain the stemness and fitness of CAR T cells in the host after infusion **(Fig. 2a)**. In order to assess the potential applicability of this approach to different CAR constructs, we chose BCMA and CD19 as established MM and lymphoma CAR targets, and ROR1 as an experimental target antigen under clinical investigation in lymphoma and solid tumors.

**Figure 2:**
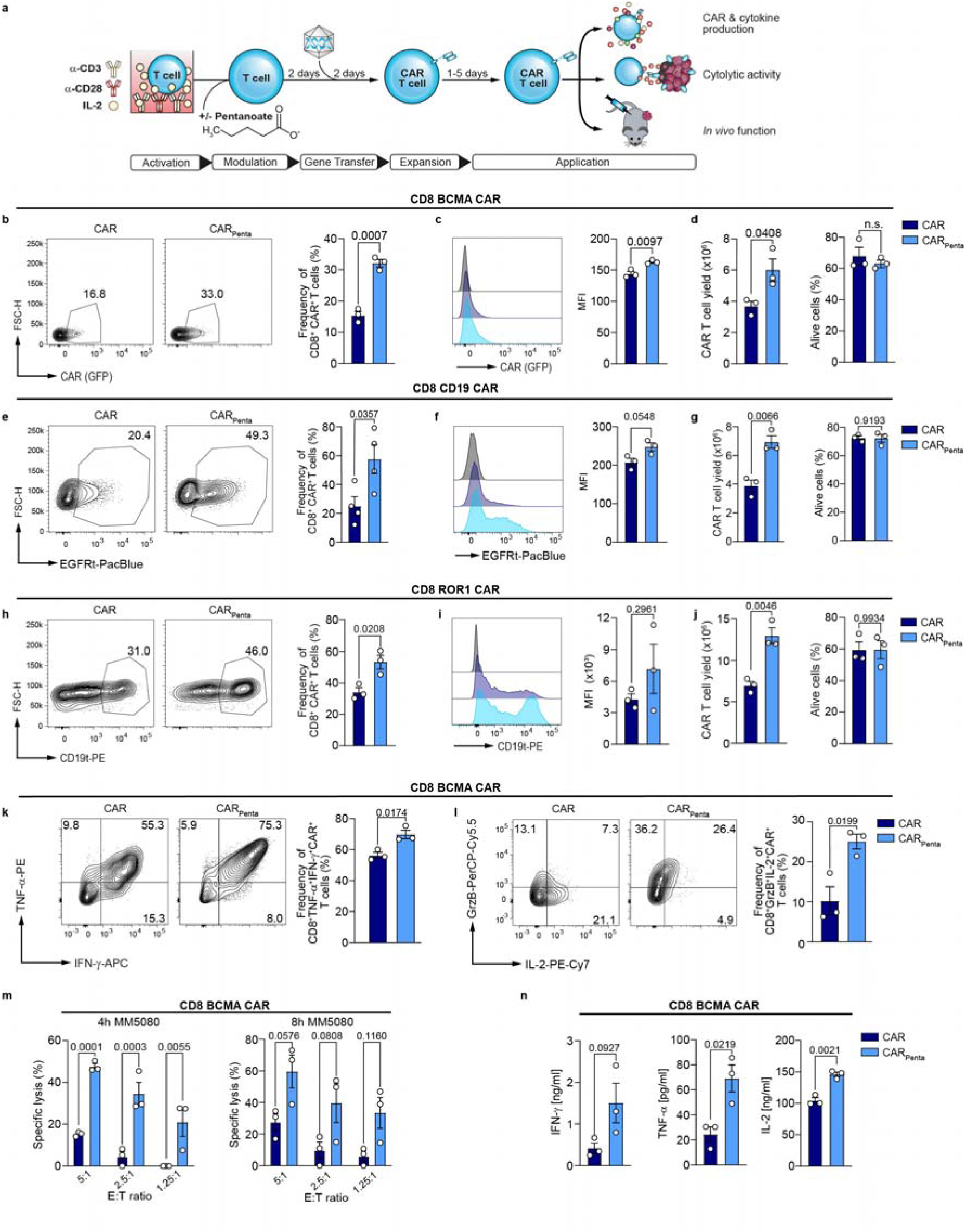
Microbial metabolites improve engineering and boost CTL-phenotype of murine CAR T cells. **a**, Schematic illustration of experimental setup for murine CAR T cell generation with pretreatment and functional analysis. **b**, Representative contour plots of the transduction marker and bar plots show frequencies of CD8^+^ BCMA- CAR T cells on day 6. Mean ± SEM from *n = 3*. **c**, Mean fluorescence intensity (MFI) of the transduction marker of generated CAR^+^ T cells. Mean ± SEM from *n = 3*. **d**, Total CAR^+^ T cell yield and viability on day 7. Mean ± SEM from *n = 3*. **e,** Representative contour plots and bar graphs show transduction efficiency of CD8 CD19- CAR T cells on day 6. Mean ± SEM from *n = 3*. **f**, Mean fluorescence intensity (MFI) for generated CAR^+^ T cells. Mean ± SEM from *n = 3*. **g**, Yield and viability of CAR^+^ T cells on day. Mean ± SEM from *n = 3*. **h**, Representative contour plots of transduction marker and bar graph show frequency of CD8^+^ ROR1- CAR T cells on day 6. Mean ± SEM from *n = 3*. **i**, Mean fluorescence intensity (MFI) of the transduction marker for generated CAR^+^ T cells. Mean ± SEM from *n = 3*. **j**, Yield and viability of ROR1**-** CAR^+^ T cells on day. Mean ± SEM from *n = 3*. **k**, Frequency of IFN-γ and TNF-α producing BCMA- CAR^+^ T cells on day 4 following antigen-independent restimulation for 5 hours. Mean ± SEM from *n = 3*. **l**, Flow cytometric analysis of Granzyme B and IL-2 production by CD8^+^ BCMA- CAR T cells after restimulation on day 4. Mean ± SEM from *n = 4*. **m**, Specific cytolytic activity of BCMA- CAR T cells at different E:T ratios at 4 and 8h timepoint. Mean ± SEM from *n = 3*. **n**, Cytokine secretion of IFN-γ, TNF-α and IL-2 after 24-hour co-culture of BCMA- specific CAR^+^ T cells with target cells. Mean ± SEM from *n = 3*. (**b**-**n**) Data represent pooled data from independent experiments. (**m** and **n**) *n=3* biological replicates; pooled data from *n=3* independent experiments; mean ± SEM was calculated for *n=3* independent experiments. Statistical analysis was performed using unpaired two-tailed Student’s *t* test (**b**-**l**) and two-way analysis of variance (ANOVA) with Tukey’s multiple-comparison test (**m** and **n**).

Isolated T cells from murine spleens and lymph nodes were activated with α-CD3 and α-CD28 antibodies in presence of pentanoate for 2 days, prior to retroviral gene transfer of the CAR transgene. Between 3 to 7 days post transduction, we characterized phenotypic and functional changes both *in vitro* and *in vivo* **(Fig. 2a)**. Generation of CD4^+^ and CD8^+^ BCMA-specific CAR T cells showed that both transduction efficacy and transduction marker expression were elevated in the pentanoate-engineered CAR T cells (CAR_Penta_) as compared to the untreated ones (CAR). Further, these features were accompanied by increased cell yield without impairment of viability **(Fig. 2b-d** and **Extended Data Fig. 1a-c)**. Similar data were obtained when the treatment was applied to the production of CD4^+^ and CD8^+^ CD19 and ROR1 CAR T cells **(Fig. 2e-j** and **Extended Data Fig. 1d-i).** Moreover, both BCMA and ROR1 CAR_Penta_ T cells showed strong upregulation of the effector cytokines TNF-α and IFN-γ, as well as granzyme B and IL-2 in an antigen-independent manner **(Fig. 2k, l** and **Extended Data Fig. 1j-o)**. Prior to the assessment of cytotoxic activity, CAR T cells were cultured for 4 days without pentanoate to explore the longevity of modulation in absence of continuous exposure. BCMA CAR_Penta_ T cells showed significant improvements in cytotoxic activity and antigen-dependent cytokine release upon co-culture with MM5080 cells as compared to conventionally engineered CAR-T cells **(Fig. 2m,n** and **Extended Data Fig. 2a,b)**. Similar effects were observed in co-cultures of ROR1-overexpressing pancreatic ductal adenocarcinoma cells Panc02 (PancROR1) and colorectal carcinoma cells MC38 (MC38ROR1) with ROR1 CAR T cells, as well as of CD19^+^ Eµ-myc cells with CD19- specific CAR-T cells **(Extended Data Fig. 2c-j)**. Interestingly, CAR_Penta_ T cells retained superior cytotoxic function when assessed 14 days after the initial pentanoate exposure (**Extended Data. Fig. 3a,b**). These results show that short-term pentanoate treatment leads to a sustained phenotype in different CAR T cell products independent from the target antigen.

### Pentanoate-engineering enhances CAR T cell efficacy, persistence and resistance to TME factors

To assess the therapeutic potential of CAR_Penta_ T cells, we established syngeneic tumor models in immunocompetent mice. We have recently described a syngeneic MM model that recapitulates the genetics and pathology of human disease. Therefore, the representation of the hostile bone marrow TME similar to the patient setting enables predictive studies on the efficacy of immunotherapy approaches [19]. Complementary, we have established a CAR targeting murine BCMA to study CAR-T cell dynamics in the syngeneic model.

For the MM *in vivo* challenge, 5×10^6^ MM5080 cells were injected i.v. into immunocompetent C57Bl/6 mice (**Fig. 3a,b**). 10 days after engraftment, 1×10^6^ BCMA CAR T cells with an equal CD4:CD8 ratio were transplanted and survival of mice monitored. While all animals treated with control BCMA CAR T cells were taken out of the experiment between day 27 and day 59 post tumor challenge, pentanoate-engineered cells mediated a significant survival advantage. These results demonstrated the superior efficacy of BCMA CAR_Penta_ T cells, in line with clinical results (**Fig. 1**).

**Figure 3:**
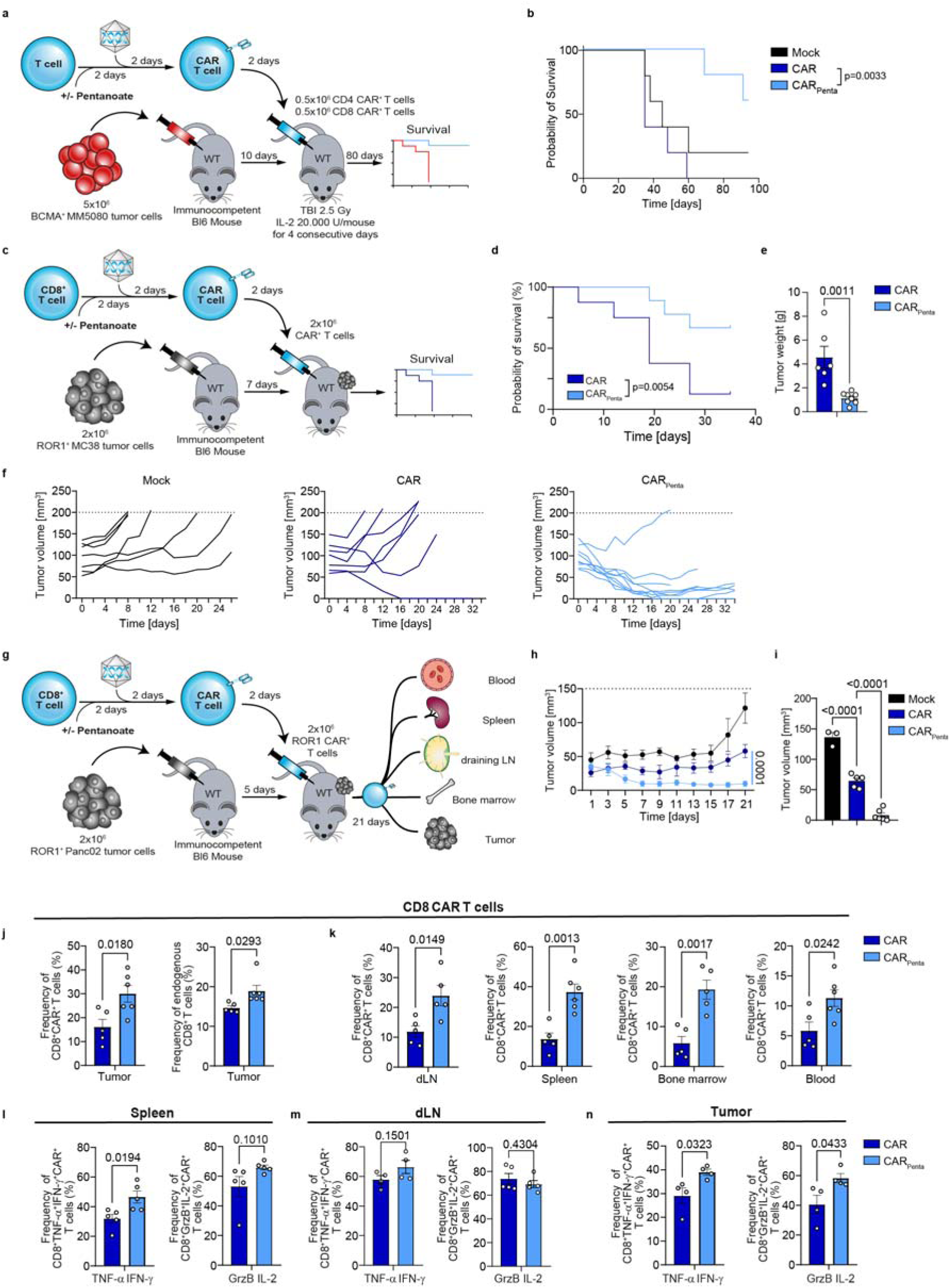
Anti-tumor reactivity in hematological and solid malignancies. **a**, Schematic representation of the experimental timeline of BCMA- CAR T cells in a MM5080 tumor model. **b**, Kaplan-Meier analysis of overall survival for all tested mice (*n* = 5 mice/group). **c**, Scheme illustrating the analysis of ROR1-CAR T cells in a MC38ROR1 tumor model. **d,** Survival of MC38ROR1 tumor bearing mice. Mean ± SEM from *n = 8* mice for control group, *n = 9* for CAR or CAR_Penta_ groups. **e**, Tumor weight at the endpoint. Mean ± SEM from *n = 6* for CAR and *n = 8* for CAR_Penta_ groups. **f**, Single tumor weight over the course of the experiment. **g**, Scheme illustrating the analysis of ROR1- CAR T cells in a PancROR1 tumor model. **h**, PancROR1 tumor growth after treatment with CD8^+^ CAR T cells. **i**, Tumor volume at endpoint day 21. Mean ± SEM from *n = 3* mice/group for mock, *n = 6* mice/group for CAR treated groups. **j**, Frequency of CD8^+^ ROR1- CAR T cells (left panel) and endogenous CD8^+^ T cells in the tumor on day 21. Mean ± SEM from *n = 5* for CAR, *n = 6* for CAR_Penta_. k, Frequency of CD8^+^ CAR T cells in the draining lymph node (dLN), spleen, bone marrow and blood on day 21. Mean ± SEM from spleen, BM and blood; Mean ± SEM from *n = 4* for CAR, *n = 5* for CAR_Penta_ for dLN. **l** – **n**, Flow cytometric analysis of IFN-γ^+^ TNF-α^+^ and GranzymeB^+^IL-2^+^-double positive ROR1-CAR T cells from the spleen (**l**), dLN (**m**) and tumor (**n**) following restimulation with PMA/Ionomycin for 5 h. Mean ± SEM from *n = 5* mice/group and *n= 4* mice/group. Survival curves were compared by the log-rank Mantel-Cox test (**b** and **d**). Statistical analysis was performed using unpaired two-tailed Student’s *t* test (**e**,**i**,**j** - **n**).

Based on these promising effects in the hostile bone marrow TME, we wanted to investigate whether the modified CAR T cells can also confer beneficial effects in different solid tumor milieus. Thus, we assessed the impact of CAR_Penta_ T cells on mouse survival in an aggressive MC38ROR1 model with hot TME characteristics (**Fig. 3c**). Monitoring over 34 days post-transfer revealed superior anti-tumor activity and survival as compared to the control group with conventionally engineered CAR T cells (**Fig. 3c-f**). While the control group only delayed tumor growth before the experimental cut-off size was reached, pentanoate-engineered CAR T cells showed much more robust tumor control.

Next, we characterized the behavior of CAR_Penta_ T cells in the Panc02 model, which develops a cold, immunosuppressive, and T cell-excluding TME. CAR_Penta_ T cells were transferred into mice bearing 5-day-old PancROR1 tumors, prior to endpoint analysis on day 21 post-transfer (**Fig. 3g**). Treatment of mice with pentanoate-engineered ROR1 CAR T cells led to significantly improved tumor control, approaching mass clearance in some mice starting 5-7 days after administration. In contrast, the conventionally engineered CAR T cell group showed inferior anti-tumor activity and trended toward loss of tumor control by the end of the experiment (**Fig. 3h** and **Extended Data Fig. 4**). Analysis at the endpoint showed low residual tumor mass and an increase in both CAR_Penta_ T cells as well as endogenous CD8^+^ tumor infiltrating lymphocytes (TILs) (**Fig. 3i,j**). Furthermore, CAR_Penta_ T cells in the draining lymph node (dLN), spleen, bone marrow and blood were more persistent (**Fig. 3k**). Intracellular cytokine staining revealed elevated frequencies of TNF-α^+^ IFN-γ^+^ and Granzyme B ^+^ IL-2 ^+^ CAR T cells in the spleen, and to a lesser extent in the dLN (**Fig 3l,m**). Significant differences were found in the tumor which reflected preservation of the cytokine profile *in vitro* (**Fig. 2h, i** and **Fig. 3n)**. Interestingly, *in vivo* transfer of a non-engineered T cell population similarly showed an antigen-independent expansion and persistence in the periphery upon pentanoate but not butyrate treatment (**Extended Data Fig. 5**). By contrast, delaying the timing of pentanoate treatment until day 2 after CAR engineering resulted in limited persistence following *in vivo* transfer into immunocompetent hosts (**Extended Data Fig. 6a-c**). This emphasizes the impact of T cell state at the time of pentanoate treatment and suggests that pentanoate-mediated programming is most efficient within an early activation window. Given the well-established HDAC-inhibitory activity of pentanoate, its impact on T cell programming may be maximized during transitional states characterized by high chromatin remodeling activity.

The enhanced tumor control by CAR_Penta_ T cells in the challenging Panc02 model suggested that pentanoate treatment could enable T cells to acquire enhanced resistance to hostile TME factors present in the solid tumors. Thus, we further characterized CAR and CAR_Penta_ T cell function in *in vitro* culture conditions designed to mimic TME-mediated T cell dysfunction.

Under hypoxic conditions, CAR_Penta_ T cells exhibited superior lytic activity, suggesting greater maintenance of effector functions despite the oxygen-reduced environment (**Extended Data Fig. 7a**). We next engineered CAR T cells under low IL-2 conditions mimicking the IL- 2-depleted environment characteristic of Treg-rich TMEs. Analysis of tumor cell survival showed that under these conditions, typically unfavorable to T cell functionality, CAR_Penta_ T cells retained superior lytic activity, potentially due to enhanced IL-2 secretion (**Fig. 2n** and **Extended Data Fig. 7b**).

Our results demonstrate that introducing pentanoate to the CAR T cell manufacturing process confers long-lasting improvements in effector function, i*n vivo* persistence and resistance to immunosuppressive factors in hot and cold TMEs, generating enhanced CAR T cell products with improved activity against hematologic and solid malignancies.

### Histone hyperacetylation enhances CAR T cell function

We next focused on elucidating the mechanisms behind pentanoate-mediated enhancement of CAR T cell function. The sustained increase in effector function after removal of the commensal metabolite suggests a lasting effect of pentanoate (**Fig. 3** and **Extended Fig. 3a,b**). This might be attributed to SCFA-mediated inhibition of histone deacetylases (HDACs) and consequently histone hyperacetylation, thereby altering the epigenetically programmed fate of eukaryotic cells (**Fig. 4a, b**). To predict how HDACs interact with fatty acids, we used AlphaFold 3 to model zinc and palmitate bound structures of class I and II HDACs (**Extended Data Fig. 8a-c**). The structural prediction revealed that HDACs of both classes coordinate palmitate in their reactive centers with high confidence up to C5 (pLDDT >90 for class I and 70-90 for class II; **Extended Data Fig. 8d).** Docking of C2-C6 SCFAs (acetate to hexanoate) into an AlphaFold 3-predicted structure of zinc cofactor-bound HDAC1 suggested that the aliphatic chains might bind in an inward facing orientation up to a chain length of C4 and in an outward facing conformation for C6 SCFAs (**Fig. 4b, c** and **Extended Data Fig. 9a)**. Notably, docking of pentanoate (C5) showed two conformations, with the best scoring inward facing conformation being preferred (predicted ΔG ≈ −8.6 kcal/mol), relative to the best outward facing orientation (predicted ΔG ≈ −7.6 kcal/mol, **Extended Data Fig. 9a)**. Docking of pentanoate into HDACs of class I and IIa suggested preferential binding to class I enzymes (**Extended Data Fig. 9b, c**). We experimentally verified the predicted inhibition of HDAC1 and 2 over class IIa by SCFAs (**Fig. 4d** and **Extended Data Fig. 9d**).

**Figure 4:**
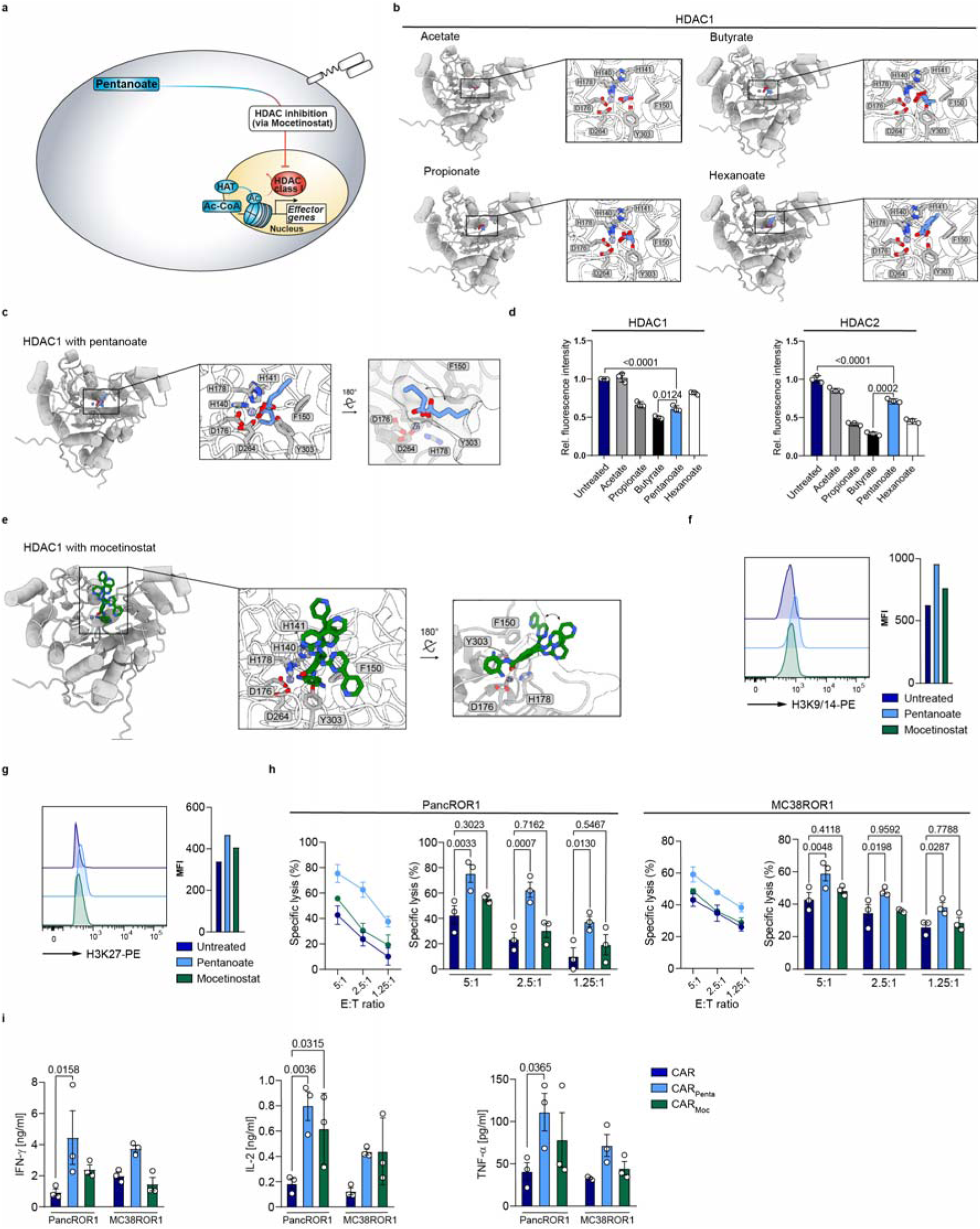
HDAC class I inhibition-mediated hyperacetylation improves CAR T cell function. **a**, Schematic pathway illustration detailing the HDAC-mediated T cell regulation. **b** and **c**, Shown are SCFAs (blue sticks), i.e. acetate, propionate, butyrate, pentanoate and hexanoate docked into AlphaFold 3-predicted structures of zinc cofactor-bound (spheres) HDAC1 (gray cartoons). **d**, Influence of bacterial SCFAs on the activity of recombinant class I and class II HDAC enzymes. **e**, Mocetinostat (blue sticks, HDAC class I inhibitor) docked into an AlphaFold 3-predicted model of zinc cofactor-bound (spheres) HDAC1 (gray cartoons). **f** and **g**, Histone acetylation status of T cells are measured by the expression of H3/K9-14 (**f**) and H3K27 (**g**) via flow cytometry treated with indicated substances. 1 out of 3 representative experiments is shown. **h**, Specific cytolytic activity of CD8^+^ CAR T cells generated in the presence of indicated substances against ROR1-expressing tumor cell lines at different E:T ratios after 6 h. Mean ± SEM from *n = 3*. **i** Cytokine secretion of CD8^+^ CAR T cells after 24 h co-culture with target antigen expressing cell lines, measured for IFN-γ, TNF-α and IL-2 by ELISA. Mean ± SEM from *n = 3*. **h** and **i**, biological replicates; pooled data from independent experiments. Statistical analysis was performed using one-way ANOVA (**h**) or two-way analysis of variance (ANOVA) with Tukey’s multiple-comparison test (**i**).

Although the SCFAs analyzed are all structurally similar and elicit HDAC inhibition, the effects of each SCFA are highly context- and cell type-dependent. Differentiation of CD4^+^ T cells towards regulatory T cells (Tregs) has long been linked to the HDAC-inhibitory activity of SCFAs [1, 16, 20]. Haradhvala and colleagues have reported that patient relapse is associated with the presence of CAR-Tregs [21]. However, only propionate and butyrate, but not pentanoate, induced the Treg master regulator Foxp3 (**Extended Data Fig. 10**). Thus, the use of pentanoate over other SCFAs in CAR-T cell engineering may reduce the risk of Treg induction in the manufactured product. Of note, treatment of inducible Tregs (iTregs) with pentanoate suppressed Foxp3 and reciprocally induced the transcription factor T-bet in a concentration-dependent manner. T-bet is a key regulator of IFN-γ in CTL and Th1 T cells which can antagonize Foxp3-mediated differentiation (**Extended Data Fig. 11a-d**). Tregs derived from T-bet-deficient (*Tbx21^-/-^*) animals were unable to express the same level of IFN-γ as WT counterparts. Moreover, comparisons between T cells generated in the presence of different SCFAs highlighted pentanoate as the most potent modulator of anti-tumor activity (**Extended Data Fig. 12a, b**). These findings suggest that the use of pentanoate could repress CAR Treg development and favor CTL/ Th1 polarization (**Extended Data Fig. 11a-d**).

To assess whether clinically relevant HDAC class I inhibitors could mimic pentanoate’s reprogramming features, we engineered ROR1 CAR T cells in presence of mocetinostat (CAR_Moc_). AlphaFold 3-modeling of zinc-bound HDAC1 and docking of mocetinostat into the reactive center suggests a binding mode similar to that of pentanoate, with a predicted ΔG of approximately −8.1 kcal/mol (**Fig. 4e** and **Extended Data Fig. 9a)**. Intracellular flow cytometry staining for histone post-translational modifications revealed an increase in acetylation within CD8^+^ and Th1 T cells at H3 Lys9/14 and H3 Lys27, respectively, both prominent marks of open chromatin and transcriptional activity (**Fig. 4f**, **g** and **Extended Data Fig. 13a**). Next, we analyzed the impact of pentanoate- and mocetinostat-mediated hyperacetylation for the CD4^+^ and CD8^+^ CAR T cell phenotype. Surprisingly, and despite mocetinostat’s strong HDAC-inhibitory activity, CAR_Moc_ T cells elicited less potent lysis of target cells and antigen-specific cytokines secretion compared to CAR_Penta_ T cells, although both were superior in killing with respect to untreated CAR T cells (**Fig. 4h**, **i**). Engineering in the presence of HDAC class II inhibitor TMP-195 did not improve CAR T cell cytotoxicity (**Extended Data Fig. 13b**), in line with our previous findings. These results confirm that epigenetic remodeling affects the CTL phenotype; however, a standalone implementation of a clinical HDACi such as mocetinostat is not sufficient to reconstitute the enhanced cytotoxicity and cytokine release of CAR_Penta_ T cells.

### Pentanoate elicits epigenetic and metabolic modulation

We further investigated whether pentanoate influences cellular metabolism and augments HDACi function (**Fig. 5a**). CD4^+^ and CD8^+^ CAR_Penta_ T cells showed increased phosphorylation of the central metabolic regulator S6, a downstream target of mTOR (**Extended Data Fig. 14a, b**). By staining with 2NBDG and BODIPY, we observed an increased glucose and fatty acid uptake (**Extended Data Fig. 14c, d)**. Moreover, CD4^+^ and CD8^+^ CAR_Penta_ T cells increased the expression of the master regulator of mitochondrial biogenesis PGC-1α and exhibited greater mitochondrial mass, compared to control CAR T cells (**Fig. 5b**, **c** and **Extended Data Fig. 14e, f**). Thus, an mTOR-mediated shift towards glucose oxidation might contribute to the CAR_Penta_ T cell phenotype. To explore this, we generated CAR T cells in presence of the clinically used drug dichloroacetate (DCA, CAR_DCA_ T cells). Of note, DCA is a pharmacological inhibitor of mitochondrial pyruvate dehydrogenase kinase (PDK) and is currently under clinical investigation for cancer therapy [22]. PDK activates pyruvate dehydrogenase (PDH), which acts as a gatekeeper enzyme for pyruvate flux into the TCA cycle. Consequently, DCA redirects metabolism from lactate fermentation to glucose oxidation in mitochondria. Similar to pentanoate, engineering of CAR_DCA_ T cells increased fatty acid uptake and mitochondrial mass (**Fig. 5c** and **Extended Data Fig. 14d**). Thus, we decided to use DCA to mimic the glucose oxidative enhancement during the engineering process.

**Figure 5:**
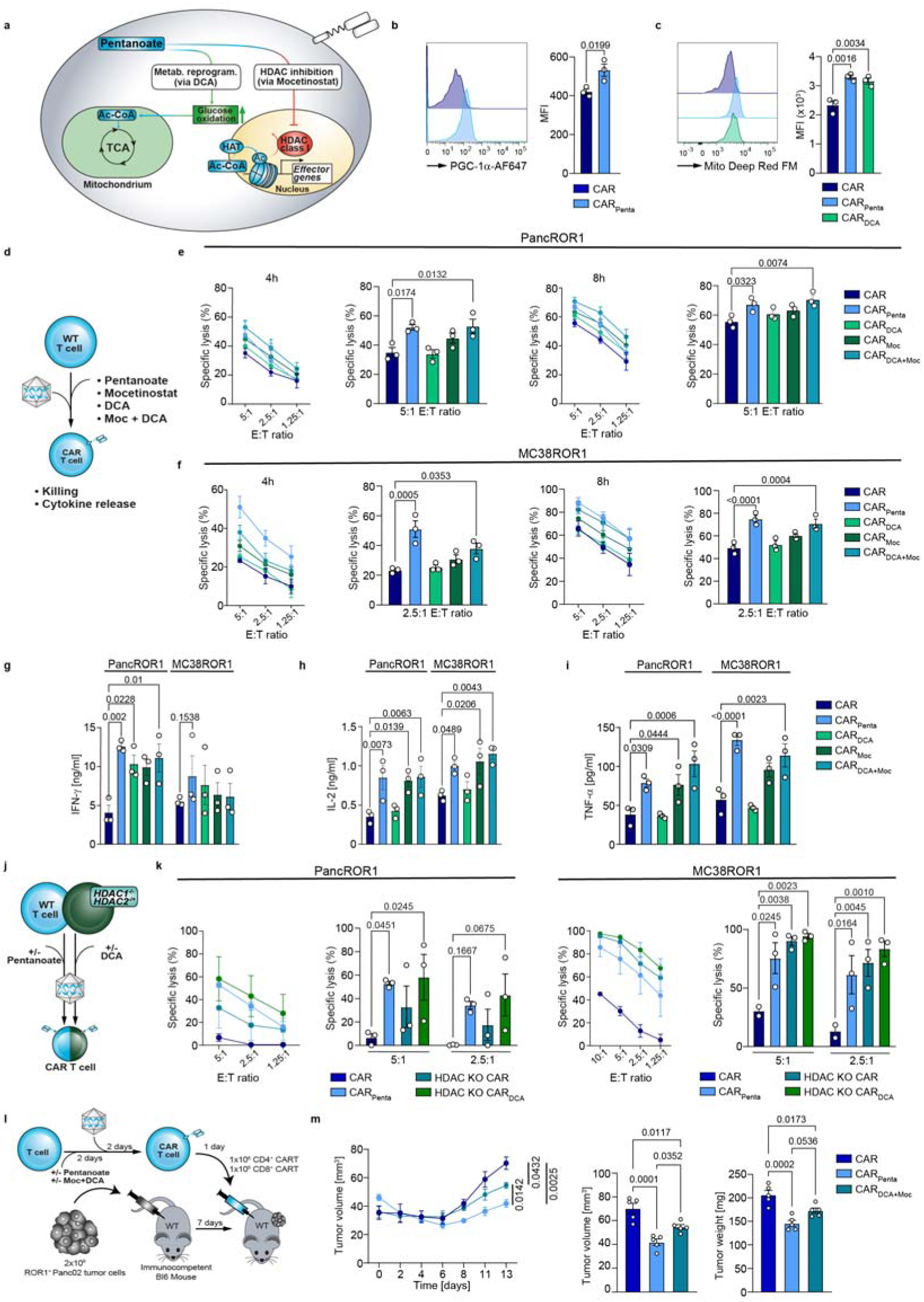
Pentanoate synergizes epigenetic and metabolic modulation to boost effector function. **a**, Schematic pathway representation. **b**, Flow cytometry analysis and quantification of PGC-1α in CD8^+^ CAR T cells either untreated or treated with pentanoate. Mean ± SEM from *n = 3*. **c**, Flow cytometric analysis of MitoFM in CD8^+^ CAR T cells treated with indicated substances. Mean ± SEM from *n = 3*. **d**, Representation of workflow with combinatorial or single treatment for CD8^+^ T cells. **e** and **f**, Cytolytic activity of ROR1- CAR T cells against PancROR1 (**e**) and MC38ROR1 (**f**) at variable E:T ratios after 4 h and 8 h. Mean ± SEM from *n = 3*. **g**-**j**, Secretion of IFN-y (**g**), IL-2 (**h**) and TNF-α (**i**) after 24 hours by CD8^+^ CAR T cells upon co-incubation with ROR1-expressing tumor cell lines. Mean ± SEM from *n = 3*. **j**, Scheme illustrating the different pretreatments of CAR^+^ T cells generated from wildtype or HDAC knockout mice. **k**, Cytolytic activity of CD8^+^ CAR T cells against target cells with different E:T ratios after 4 h. CAR T cells were generated as shown in **j**. Mean ± SEM from *n = 3*. **l**, Experimental setup of pretreatment with pentanoate or DCA/mocetinostat combination during the CAR^+^ T cell manufacturing process followed by injection in PancROR1 tumor bearing mice. **m**, Tumor growth (left) over time as well as tumor volume (middle) and tumor weight on day 14 (right) after tumor inoculation are shown. Mean ± SEM from *n = 10* mice/group until day 7, then *n = 5* mice/group until day 14. **b, c, e**-**l** and **k**, independent experiments. **e**-**l**, **k**, biological replicates; pooled data from independent experiments. Statistical analysis was performed using one-way ANOVA (**b**, **c**, **m**) or two-way analysis of variance (ANOVA) with Tukey’s multiple-comparison test (**e**-**I** and **k**).

Next, we tested whether potential synergy between HDAC inhibition and metabolic reprogramming could explain superior CAR_Penta_ T cell effector function. To do so, we engineered CAR T cells in the absence or presence of pentanoate, DCA, mocetinostat or a combination of the latter, respectively (**Fig. 5d**). These were subjected to co-cultures using PancROR1 or MC38ROR1 tumor cells to assess cytotoxicity (**Fig. 5e**, **f**). Interestingly, while CAR_DCA_ T cells did not show a significant improvement in killing, combined administration of DCA and mocetinostat during manufacturing (CAR_DCA+Moc_ T cells) resulted in an additive effect which was also reflected in antigen-dependent secretion of TNF-α, IFN-γ and IL-2 (**Fig. 5g-i**). Since the co-stimulatory domain within CAR constructs has been shown to alter cellular metabolism and therefore CAR T cell function [23], we performed a set of experiments using a CD28-based ROR1 CAR. This resulted in similar effects upon treatment with the previously mentioned drugs with regards to cytotoxicity and mitochondrial mass (**Extended Data Fig. 15a, b**).

To understand a potential contribution of mocetinostat off-target effects, we tested HDAC class I involvement by using murine T cells with homozygous HDAC1 (*HDAC1*^-/-^)- and heterozygous HDAC2 (*HDAC1*^-/-^)- deficiency **(Fig. 5j**). HDAC-deficient CAR T cells showed superior specific lysis compared to the WT CAR control, mimicking the trends observed in CAR_Moc_ T cells **(Fig. 5k**). Engineering in the presence of DCA further boosted lytic activity and cytokine secretion, comparable to levels of CAR_Penta_ T cells (**Fig. 5k** and **Extended Data Fig. 16)**. This demonstrates a synergistic relationship between HDAC inhibition and metabolic enhancement that contributes to the CAR_Penta_ T cell phenotype.

To evaluate whether the *in vitro* results could be recapitulated *in vivo*, we applied suboptimal doses of CAR, CAR_Penta_ and CAR_DCA+Moc_ T cells in the Panc02ROR1 model characterized by a cold TME, without lymphodepletion (**Fig. 5l)**. Surprisingly, differences in tumor mass and a lack of peripheral expansion indicated that the conventional drugs were not able to reproduce the efficacy of pentanoate, although improved tumor control was observed in both groups as compared to the control CAR T cells (**Fig. 5m** and **Extended Data Fig. 17a, b)**. Thus, pentanoate involves unique mechanisms that are not affected by mocetinostat and DCA.

### Pentanoate is metabolized in the TCA and becomes part of the epigenetic imprint

While DCA and pentanoate both influence mitochondrial activity, they might differ in their effect on glycolytic and oxidative pathway intermediates (**Fig. 6a**). To trace the pathway intermediates, we pulsed CAR T cells with either ^13^C-glucose or -glutamine tracers, prior to analysis via GC-MS (**Fig. 6b**). Tracing of ^13^C-glucose revealed that less glucose was metabolized to lactate upon DCA treatment, confirming its activity as PDK inhibitor. No significant changes in pyruvate were observed for any of the small molecules used (**Fig. 6c**). Of note, we detected a strong decrease in glucose-derived citrate in CAR_Penta_ T cells that was not observed for mocetinostat or DCA (**Fig. 6c**). The impact on glucose-derived citrate was shown by its reduction of approximately 25% as compared to control CAR T cells. This trend was more pronounced with a reduction of approximately 50% labeling for downstream TCA intermediates such as α-ketoglutarate, fumarate and malate. In contrast, although DCA reduced cellular lactate levels, its effects on glucose oxidation did not change the overall glucose-dependent citrate production. These data highlight pentanoate’s capacity to reprogram the TCA in T cells by modulating the generation of glucose-derived citrate contrasting with conventional drugs.

**Figure 6:**
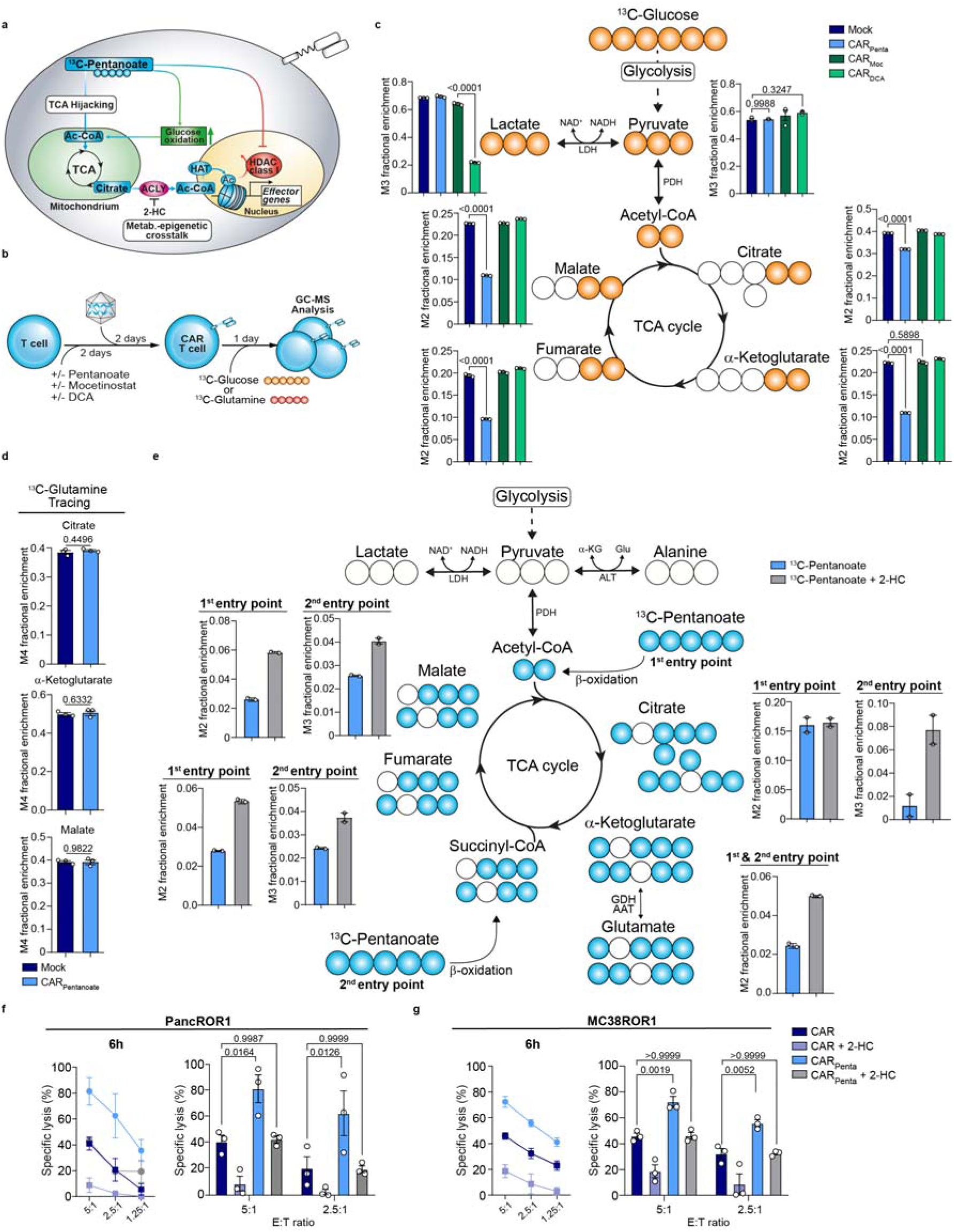
Pentanoate hijacks the cellular metabolism and becomes part of the epigenetic imprint. **a**, Scheme illustrating pentanoate modulation and metabolism hijacking of T cells. **b**, Schematic for the generation of CAR^+^ T cells in combination with pretreatment prior to metabolite tracing **c**, GC-MS isotope tracing of ^13^C-glucose-derived metabolites in CAR T cells engineered in the presence of specified substances. Histograms show fractional enrichment of ^13^C-glucose-derived TCA metabolites. Mean ± SEM from *n = 3* biological replicates. **d**, GC-MS isotope tracing of ^13^C-labeled glutamine in untreated or pentanoate-treated CAR T cells regarding citrate, α-ketoglutarate and malate. Mean ± SEM from *n = 3* biological replicates. **e**, GC-MS tracing of ^13^C-pentanoate in CD8^+^ CAR T cells pretreated with indicated substrates. Histograms show the frequency of pentanoate-derived carbons in the downstream TCA cycle intermediates. *n=2* independent T cell pools. **f** and **g,** PancROR1 (**f**) or MC38ROR1 (**g**) tumor killing of CAR^+^ T at different E:T ratios after 6 h. Mean ± SEM from *n = 3*. Statistical analysis was performed using unpaired two-tailed Student’s *t* test (**d**), one-way ANOVA (**c**) or two-way analysis of variance (ANOVA) with Tukey’s multiple-comparison test (**f** and **g**).

We next addressed which alternative source was used by CAR_Penta_ T cells to replenish the citrate carbons. Upon butyrate treatment, T cells may fuel the TCA via glutamine anaplerosis to uncouple it from glycolysis [24]. By contrast, pulsing of CAR_Penta_ T cells with ^13^C-glutamine did not show a preference for incorporation into the TCA intermediates (**Fig. 6d**). Hence, fatty acid oxidation and metabolization of pentanoate might contribute to citrate generation. To understand how pentanoate is metabolized, we administered ^13^C-pentanoate to the CAR T cell culture. GC-MS analysis revealed the incorporation of pentanoate-derived carbons into citrate via the acetyl-CoA entry point. Additionally, the labeled carbons entered the TCA through a second entry point via the succinyl-CoA route (**Fig. 6e**). This is a unique feature of pentanoate attributed to the C5 aliphatic chain, which cannot be emulated by acetate, propionate or butyrate. The combined citrate amounts derived from both entry points correspond to the gap detected in the previous glucose tracing experiment which underlined a limited preference for pentanoate over glucose (**Fig. 6c, e**). Besides TCA intermediates being required for energy homeostasis, conversion of citrate into acetyl-CoA by the nuclear ATP-citrate lyase (ACLY) serves as a source for histone acetylation. This process links metabolism and epigenetic regulation [25]. To test whether ACLY directs the pentanoate-derived citrate flux, we inhibited ACLY via 2-hydroxy citrate (2-HC) in the presence of the ^13^C-pentanoate tracer (**Fig. 6e**). Interestingly, 2-HC treatment of CAR_Penta_ T cell not only led to a 2-fold increase in pentanoate labeling of glutamate, fumarate and malate downstream of citrate from the first entry point, but to an additional 4-fold increase of pentanoate-derived citrate from the second one. This suggests that the latter might be used for histone acetylation in the nucleus and has accumulated upon ACLY blockade in the TCA. To investigate the functional outcome of this crosstalk, we evaluated CAR_Penta_ T and control CAR T cells that were engineered in the absence and presence of 2-HC in killing assays. Specific lysis and antigen-specific cytokine analysis showed that 2-HC administration reduced the functionality of CAR_Penta_ T cell significantly (**Fig. 6f**, **g** and **Extended Data Fig. 18a-c**).

These results demonstrate that the unique metabolization of pentanoate contributes to the cellular citrate pool that is used to bridge TCA and chromatin remodeling, emphasizing the importance of the epigenetic-metabolic crosstalk for effector T cell function.

### Pentanoate-engineered T cells elicit a naïve-like phenotype in the TME

To better understand the effects of the drug treatments on the immune microenvironment at, we performed single cell RNA sequencing (scRNA-seq). E*x vivo* isolated immune cells were obtained from tumor-bearing mice treated with CAR, CAR_DCA+Moc_ and CAR_Penta_ T cells on day 7 and day 14 post transfer, respectively, and the pre-infusion products prior to scRNA-seq (n=24, n=4 mice /treatment/ point in time, **Fig. 9a**). After performing the QC steps a total of 16908 cells were identified. The UMAP representing the distribution of the cells based on conditions, time points and mouse of origin are presented in **Extended Data Fig. 19a-c**. By using scGate we were able to annotate these cells and identify 4774 of pure T cells that were further divided in 14 clusters based on their cell state (**Figure 7b, c** and **Extended Data Fig. 19d**). Of interest, following treatment with pentanoate we observed an enrichment of CD4 and CD8^+^ naïve-like T cells (TCF7^+^, CCR7^+^, PD1^-^, TOX^-^) while a decrease in CD8^+^ TEX (GZMK^-^, TCF7^-^, TOX^+^, HAVCR2^+^, PD1^+^) was noted (**Figure 7d,e**).

**Figure 7:**
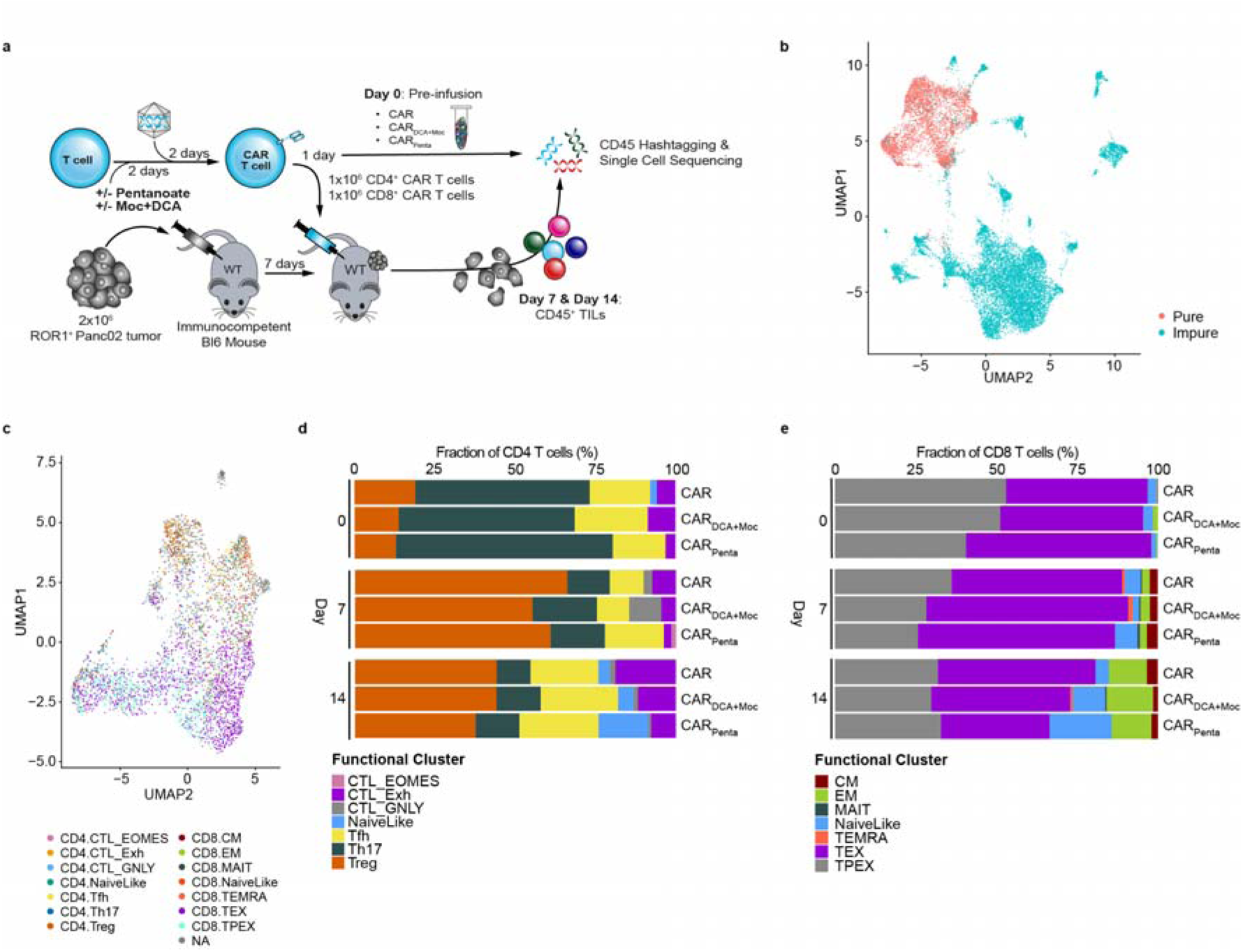
Epigenetic-metabolic reprogramming induces a naïve-like phenotype *in vivo*. **a,** Schematic representation of PancROR1 tumor model for the collection of T cells for scRNA seq. **b**, Uniform Manifold Approximation and Projection (UMAP) showing the distribution of immune cells collected from mice using scRNAseq data in the LSI space. Each point represents one cell. The cells are marked by color code based on cell annotations. Red indicates T cells, blue non-T cells. **c**, UMAP showing the distribution of T cells based on their cell states. The cells are marked by color code based on the different cluster they belong to. **d** and **e**, Barplots indicating the changes in percentage of the different CD4 (**d**) and CD8 (**e**) T cell subsets noticed following the different treatments and time points (day 0, 7 and 14).

Taken together, these data provide evidence for pentanoate’s ability to increase the fitness of T cells, create a more active immune microenvironment that supports anti-tumor activity and reduce T cell exhaustion.

## Discussion

Both homeostatic regulation and pathophysiologic development have been linked to the composition of the gut microbiome. The direct interaction between host and commensals, as well as postbiotics reaching distant sites in the body, are major influences on the immune system which can be therapeutically exploited. Here we demonstrate that the microbial metabolite pentanoate is predictive for patient survival and can be integrated into the CAR T cell life cycle to exploit the benefits of the host-microbiome interaction.

Previous reports in the context of ICI have revealed that commensals such as *Bifidobacterium* and *Akkermansia muciniphila* enhance anti-tumor immunity in mice in a T cell-dependent manner and are associated with better patient prognosis [4, 5, 8]. In line, correlations between antibiotics treatment and worse patient outcomes were recently established for CAR T cell therapy (**Fig. 1a**, **b**) [9, 8]. Initial attempts to exploit microbiome-based approaches for cancer immunotherapy involved fecal microbiota transplantations from responders into ICI-resistant patients after microbiome reset as well as patients with severe gastrointestinal graft-versus-host disease (GVHD) following allogeneic stem cell transplantation (allo-SCT) [26, 27]. A smaller fraction of the ICI-treated patients indeed achieved a response, but the remainder of the cohort remained resistant. Likewise, the overall response rates of GVHD patients were between 30 and 50% [28].

Furthermore, contradictory findings have slowed the translation of other microbiome-based therapies. Similar to our findings in CAR T patients, detection of high levels of the SCFAs propionate and butyrate was correlated with decreased PFS in ICI patients (**Fig. 1d**) [15]. The same class of metabolites was initially described as potent gatekeeper of intestinal homeostasis capable of inducing Treg differentiation and repressing allergic, inflammatory and autoimmune reactions [20, 16, 10, 29]. By contrast, we have demonstrated that SCFAs induce not only regulatory T and B cells but also CTL-associated factors in CD8^+^ T cells **(Fig. 2)** [10]. This emphasizes cell type- and context-dependent effects of commensal metabolites rather than a broadly applicable phenotypic change [29, 17, 30, 24]. As many modes of action shaping immunosuppressive and –stimulatory phenotypes overlap, an in-depth mechanistic understanding of each commensal metabolite is required to unlock their full potential for the development of new therapeutic approaches such as CAR T cell therapy. Here, we show that among all SCFAs, pentanoate is the only one predictive of patient survival in a three-centric German CAR T cell cohort (**Fig. 1a-d**). Despite its HDAC-inhibitory function, which is a shared feature among SCFAs, pentanoate did not induce Treg differentiation in CD4 T cells in contrast to butyrate and propionate (**Extended Data Fig. 10a, b)**. Notably, the presence of CAR Tregs in infusion products can have detrimental effects in respect to relapse in patients [21]. In line with our *in vitro* findings, a reduction of Tregs in the pre-infusion product (D0) and among the TILs (D14) was observed in the CAR_Penta_ T cell group (**Fig. 7d)**. Thus, the choice of an HDAC inhibitor must be carefully made based on its interaction with the raw material when engineering immune cells. Furthermore, the importance of epigenetic regulation has been highlighted by the generation of mice bearing HDAC class I-deficient T cells whose CD4 T cells commit to the CD8^+^ lineage during aging [31]. Additionally, other epigenetic modifications such as methylation were shown to be factors determining the fate and function of CAR T cell products [32, 33].

Besides being natural HDAC inhibitors, SCFAs modulate cellular metabolism. The link between metabolic fitness and potency of cell products has prompted approaches to actively shape metabolic pathways by overexpression of nutritional transporters or master regulators like PGC-1α [34–37]. In particular, mitochondrial metabolism has been identified as a critical factor in T cell memory formation [34, 38, 24, 39]. The choice of the CAR costimulatory domain was suggested to affect efficacy and persistence of the engineered T cell via balancing of glycolytic and mitochondrial function [40, 23]. In agreement with this idea, pharmacologic inhibition of the mitochondrial pyruvate carrier in CAR T cells results in improved anti-tumor response [38]. However, we demonstrated that DCA did not improve CAR T cell efficacy to the same extent as pentanoate, despite engaging similar pathways. In contrast to clinically available drugs, pentanoate entered the TCA through two different entry points post β- oxidation, which is a unique feature of the odd-numbered C5 aliphatic chain. Of note, Bachem and colleagues reported that pentanoate’s structurally closest relative butyrate elicits its metabolic function primarily via shift towards glutamine anaplerosis [24].

The enzyme ACLY links metabolic output and histone acetylation in mammalian cells [25]. Its inhibition via 2-HC in pentanoate-engineered CAR T cells led to accumulation of SCFA-derived citrate in the TCA, primarily from the succinyl-CoA entry route, and consequently a decrease of cytotoxic function (**Fig. 6e-g**). These findings suggest that pentanoate-derived carbons contribute to histone acetylation and differentiation towards an effector phenotype.

Strikingly, superior anti-tumor activity was achieved upon a single treatment with pentanoate and retained after removal, both *in vitro* and *in vivo* (**Fig. 3** and **Extended Fig. 3**). CAR_Penta_ T cell administration resulted in enhanced tumor control compared to conventionally engineered T cells in three different syngeneic, immunocompetent models with cold and hot TME derived from hematologic and solid tumor entities, respectively (**Fig. 3**). Short-term pentanoate treatment led to enhanced expansion and persistence in the periphery, greater tumor infiltration and a naive-like differentiation with a decrease in exhaustion (**Fig. 3j-k**). These observations suggest a broadly applicable phenotype that might be used especially to overcome the limitations of CAR T cell therapy in solid tumors. By contrast, the effect of butyrate on T cells in the context of infection was previously reported to induce a memory phenotype [24]. Pentanoate-engineered CAR T cells maintained improved lytic activity under hypoxia and low IL-2 conditions suggesting resistance against TME-mediated dysfunction (**Extended Data Fig. 7**). Moreover, the enhanced autocrine IL-2 secretion may facilitate competitive fitness over endogenous immune cells, even in absence of lymphodepletion. While administration of lymphodepleting drugs is typically a prerequisite to generate a niche for CAR T cells, it renders patients susceptible to infections and necessitates preventive antibiotic treatment. This countermeasure in turn can affect pentanoate abundance and survival upon CAR T cell treatment (**Fig. 1a-d**) [9, 8]. Thus, approaches requiring minimal lymphodepletion with retained immune cell activity would be favorable for clinical application.

Although previous studies highlighted the relevance and potential of microbiome modulation in influencing immunotherapy responses, the clinical implementation is only slowly picking up pace. Designer consortia, pre- and antibiotics or phages face challenges as the dynamics of commensal colonization and the diversity of host immune factors might complicate their application [41, 6, 30]. The use of probiotic strains bears the risk for translocation and infections as well as causing adverse effects due to their spectrum of postbiotics. Our study introduces postbiotic pentanoate as a promising model of the microbiome-host relationship, highlighting its advantages by incorporating epigenetic and metabolic adjustments into CAR T cell products, independently of gene editing, CAR target and co-stimulatory domain (**Fig. 2b-m**, **Extended Data Fig. 1** and **Extended Data Fig. 15a, b**). This approach enables a safer and more manageable side effect profile while reducing the complexity and genotoxicity associated with multi-factor genetic engineering. The integration of microbial metabolites in clinical-grade GMP manufacturing processes can offer prolonged modulation, improved metabolic fitness and standardized medicine development. Pentanoate exemplified that postbiotics are actively incorporated into immune cell metabolism, become part of the epigenetic landscape and hence contribute to determining cell function and fate. This highlights a new mechanism of host-microbe communication [42]. Its unique characteristics and predictive potential emphasize the microbiome as a source to mine for novel physiological drugs and biomarkers that can be exploited in tailored immunotherapy settings.

## Methods

### Study design and patients

The study was designed as a prospective observational study. All patients receiving CAR T cell therapy were eligible for this study regardless of the target antigen. Participants were enrolled from 05/2020 to 12/2023 at two centers: the University Medical Center Regensburg and the University Hospital Heidelberg. All patients provided informed written consent. The study was approved by the Institutional Ethics Review Board of the University of Regensburg, vote no. 21-2521-101. Patient characteristics are summarized in **Suppl. Table 1**. Fecal samples were collected from all patients prior to CAR T cell infusion. Fecal samples were prepared and short-chain fatty acids were measured as previously described [43]. Briefly, metabolites from 100mg feces were extracted using a 15-ml bead-beater tube (CKMix50, 15 ml, Bertin Technologies) filled with 2.8-mm and 5.0-mm ceramic beads and a bead-beater (Precellys Evolution, Bertin Technologies) at 10,000 rpm (3 rounds of 30s with 15s breaks). Methanol-based dehydrocholic acid extraction solvent (5 ml, c = 1.3 μmol l−1) was added as an internal standard to account for work-up losses. Short chain fatty acids were measured using the 3-NPH method in a QTRAP 5500 triple quadrupole mass spectrometer (SCIEX) coupled to an ExionLC AD (SCIEX) ultra-highperformance liquid chromatography system. Data were analyzed with MultiQuant 3.0.3 (SCIEX) and MetaboAnalyst. Stool samples have limited availability due to the individual amount obtained from each patient.

An external cohort of 60 patients with large B cell lymphoma or follicular lymphoma, treated with commercial CD19-directed CAR-T cells, was used to validate the association between stool pentanoate levels and CAR T therapy outcomes (**Suppl. Table 2**). Stool samples collected between day −30 and day 0 (median: day −3 [IQR: −4.3 to −1]) were aliquoted and stored at −80°C. One sample per patient was sent to Metabolon for metabolomic profiling

Sample Preparation: Feces samples (fresh/frozen) are analyzed for eight short chain fatty acids: acetic acid (C2), propionic acid (C3), isobutyric acid (C4), butyric acid (C4), 2-methyl-butyric acid (C5), isovaleric acid (C5), valeric acid (C5) and caproic acid (hexanoic acid, C6) by LC-MS/MS. Samples are spiked with stable labelled internal standards and are homogenized and subjected to protein precipitation with an organic solvent. After centrifugation, an aliquot of the supernatant is derivatized. The reaction mixture is diluted, and an aliquot is injected onto an Agilent 1290 Infinity or Infinity II/ Sciex QTrap 5500 or 6500 LC MS/MS system equipped with a C18 reversed phase UHPLC column. The mass spectrometer is operated in negative mode using electrospray ionization (ESI).

Sample Analysis: The peak area of the individual analyte product ions is measured against the peak area of the product ions of the corresponding internal standards. Quantitation is performed using a weighted linear least squares regression analysis generated from fortified calibration standards prepared immediately prior to each run. LC-MS/MS raw data are collected and processed using AB SCIEX software Analyst 1.6.3 and processed using SCIEX OS-MQ software

QA/QC: Three levels of QCs are prepared in feces by diluting and/or spiking with stock solutions to obtain the appropriate concentrations for each level (low/med/high). Accuracy will be evaluated using the corresponding QC replicates in the sample runs. Targeted acceptance criteria are at least 50% of QC samples at each concentration level per analyte should be within ±20.0% of a set mean and at least 2/3 of all QC samples per analyte should fall within ±20.0% of the corresponding mean.

### Animals

C57BL/6 wild-type mice were purchased from Charles River and maintained under specific pathogen free conditions at the Center for Experimental Medicine (ZEMM) at the University of Würzburg. Tbx21^−/−^ □mice (on C57BL/6 background) were maintained under specific pathogen free (SPF) conditions at the animal facility of the Philipps-University of Marburg, Germany. Mice were kept under a 12/12 h light/dark cycle between 20-24 °C in individually ventilated cages. Mice had access to standard chow and autoclaved water ad libitum and the health status of the animals was inspected by responsible animal caretakers. Male and female mice between 8-12 weeks old at the time of the experiment were used in this study. All animal protocols were approved by government (Approval number: 1457, Regierung von Unterfranken, Bayern, Germany).

Hdac1^fl/fl^Hdac2^fl/fl^ (HDAC1-2^cKO^) CD4-Cre mice (Mouse Genome Informatics [MGI] 4440556 for Hdac1; MGI 4440560 for Hdac2) were previously described and kept under specific pathogen free conditions at the Medical University of Vienna [31, 44].

### Cell lines

The mouse MC38 colon adenocarcinoma cell line was provided by the lab of Tobias Bopp/Toska Bohn. The mouse Panc02 OVA pancreatic tumor cell line was gifted from the Christian Bauer lab. All solid tumor cell lines were cultured in standard DMEM (Gibco) supplemented with 10 % heat-inactivated FCS (Gibco) and 1 % penicillin/streptomycin (Gibco) at 37 °C with 5 % CO2.

The mouse lymphoma cell line E-myc was supplied by the lab of Dirk Busch and cultured in standard RPMI 1640 (Gibco) with heat-inactivated 10 % FCS (Gibco) and 1 % penicillin/streptomycin (Gibco) at 37 °C with 5 % CO2.

The Platinum-E retroviral packaging cell line (Cell Biolabs) was cultured in standard DMEM (Gibco) supplemented with 10 % heat-inactivated FCS and 1 % penicillin/streptomycin (both Gibco) at 37 °C with 5 % CO2. Following thawing, PlatinumE cells were selected for 5 days using 1 µg/ml puromycin and 10 µg/ml blasticidin S (both Invivogen) and subsequently cultured. All cell lines were tested for mycoplasma contamination.

### CAR constructs

The murine ROR1 (R11)-CAR construct in a Vh-Vl direction with a (G4S)3-linker comprises an IgG4-hinge domain, a murine CD28 transmembrane domain and an intracellular murine 41BB followed by murine CD3zeta domain. The CAR scFv and the murine truncated CD19 transduction marker are separated via a P2A sequence. For some experiments, the 41BB costimulatory domain was replaced by a CD28 costimulatory domain. The murine CD19- CAR was designed in Vh-Vl direction with a CD8a spacer and transmembrane domain followed by CD28 co-stimulatory domain and a CD3zeta domain. Downstream of the CAR; the construct encoded a human truncated EGFR as a transduction marker separated by a P2A sequence. All vectors were cloned into a MP71-backbone.

The murine BCMA CAR was constructed using an anti-murine BCMA single-chain variable fragment (scFv) composed of the VH chain linked to the VL chain by a GGGSx4 spacer. This scFv was linked to murine CD8 spacer and transmembrane domains, followed by the 4-1BB co-stimulatory domain and the CD3ζ signaling domain. This cassette was cloned downstream of a GFP-P2A sequence in a pRubiG backbone.

### Virus production

Platinum-E cells for retroviral packaging were co-transfected with the desired construct in addition with retroviral packaging construct pCL-10A1, using the Effectene transfection reagent (QIAGEN) according to the manufacturer’s instructions. The retroviral supernatant was collected 48 and 72 h after transfection, pooled and stored at −80 °C until use.

### *In vitro* T cell differentiation and culture

For *in vitro* cultures, murine T cells from male or female mice were isolated from single cell suspension of spleen and lymph nodes. Isolation of CD8^+^ T cells was performed by positive selection (Miltenyi Biotech), followed by a negative selection for CD4^+^ T cells (Invitrogen). T cells were cultured unless stated otherwise in modified RPMI 1640 medium (Gibco) supplemented with 10 % heat-inactivated FCS (Gibco), 50 µM 2-mercaptoethanol (Gibco), 1 % penicillin/streptomycin and 1 % GlutaMAX-I (Gibco) at 37 °C with 5 % CO_2_.

Culture plates were pre-coated with 10 µg/ml polyclonal anti-hamster IgG (MP Biomedicals) for 2 h and washed once with PBS (Gibco). For Th1 differentiation, 0.5×10^6^ CD4^+^ T cells were activated in the presence of 1 µg/ml anti-CD3 (Biolegend, 145-2C11), 1 µg/ml anti-CD28 (Biolegend, 37.51), 1 µg/ml anti-IL-4 (Biolegend, BVD4-1D11), 50 U/ml recombinant human (rh) IL-2 (Miltenyi Biotech) and 10 ng/ml IL-12 (Miltenyi Biotech).

For supoptimal CTL differentiation, 0.5×10^6^ CD8^+^ T cells were cultured with 1 µg/ml anti-CD3, 1 µg/ml anti-CD28, 1 µg/ml anti-IFN-γ (Invitrogen, XMG1.2), 1 µg/ml anti-IL-4 and 50 U/ml rh IL-2.

For iTreg differentiation, CD4^+^ T cells were activated with 1 µg/ml anti-CD3, 0.5 µg/ml anti-CD28, 100U/ml rh IL-2, 2 µg/ml anti-IFN-γ, anti-IL-4 and 2 ng/ml rhTGF-β1 (R&D Systems).

In some experiments, medium of cells was added with 2 mM sodium Pentanoate (Ambeed), 5 mM dichloroacetate (Merck), 100 nM mocetinostat (Biomol), 1 µM TMP-195 (Biomol), 1 mM Butyrate (Sigma), 2 mM Propionate (Merck), 10 mM Acetate (Merck) or a combination of DCA and mocetinostat once 2 hours after activation. Furthermore, 5 mM 2-hydrocycitrate (2-HC) (Sigma) was added to cell culture at indicated time points. All exogenous metabolites were dissolved in DMSO solutions according to the manufacturer’s recommendation.

### Generation of murine CAR T cells

48 hours after activation, the medium of the cells was removed and stored at 4°C. Cells were transduced with retroviral supernatant in addition with 10 µg/ml polybrene by spin-infection at 800 rpm for 2 hours at 32 °C and placed in an incubator for 4 hours. Afterwards, the supernatant was replaced by the stored medium. The next day, T cells were put into new culture-plates. On day 4, fresh medium including 50 U/ml rh IL-2, 10 ng/ml IL-7 and 10 ng/ml IL-15 (all Miltenyi) was added. Cells were maintained between 0.5×10^6^ and 2×10^6^ cells/ml and expanded until day 7.

### Flow cytometry

T cells were harvested, washed once with PBS and stained with live/dead marker (Biolegend) for 20 min at RT. After another washing step, surface antigens with fluorophore-conjugated antibodies were stained for 20 min in PBS at 4 °C in the dark. For staining of intracellular cytokines, cells were stimulated with 100 ng/ml phorbol-12-myristat-acetate (PMA) and 1 µg/ml Ionomycin in the presence of 5 µg/ml brefeldin A for 4-5 hours at 37 °C. Cells were fixed with 2 % paraformaldehyde (Invitrogen) for 20 min at 4°C, before being stained for intracellular cytokines with antibodies in Saponinbuffer for 30 min at 4 °C.

For the detection of transcription factors and histone modifications, activated T cells were fixed with the Foxp3 / Transcription Factor Staining buffer set following manufacturer’s instructions (eBioscience) and staining with described antibodies was conducted for 30 min in Saponinbuffer at 4 °C. To assess neutral lipid content, cells were stained with BODIPY 493/503 (Invitrogen) for 20 min at RT. For the quantification of mitochondrial volume, T cells were labeled with 200 nM MitoTracker Deep Red (Invitrogen) and stained for 30 min at 37 °C. To measure glucose uptake, cells were incubated with fluorescent glucose analog 2- NBDG (Cayman Chemicals) for 20 min at 37 °C. To assess phosphorylation of intracellular S6 cells were stained as described previously [18].

All data was collected on a BD Canto II and analyzed using FlowJo. MFI marks median fluorescence intensity calculated by FlowJo. For the list of antibodies used for flow cytometry see **Suppl. Table 2**. Examples of the gating strategy are shown in **Extended Data Fig. 20**.

### Cytotoxicity assay

To determine antigen-specific tumor cell lysis, 5 × 10^3^ ffLuc-transduced tumor cells were co-cultured with CAR-transduced or untransduced (UTD) T cells at effector:target (E:T)-ratios of 10:1, 5:1, 2.5:1 or 1.25:1 in triplicates in RPMI medium supplemented with 150 ng/ml D-Luciferin. For experiments under hypoxia conditions, cells were plated in an incubator with 2% O_2._ The bioluminescence signal was measured on a Tecan Infinite 200 PRO plate reader after 4, 6, 8 and 24 h. Specific lysis was determined in reference to the corresponding untransduced T cells.

### Cytokine secretion

Secretion of cytokines for murine CAR T cells was detected by ELISA kits for mouse IFN-γ, IL-2, Granzyme B (all Biolegend) and TNF-α (Invitrogen) according to the manufacturerer’s instructions. Cells were co-incubated with 5:1 or 2.5:1 E:T ratio with antigen-presenting tumor cells in triplicates for 24 h. Absorbance was measured using a Tecan Infinite 200 PRO plate reader.

### HDAC docking methods

Structure predictions of HDACs 1-5, 7-9 with zinc ions (one zinc for HDACs 1 and 2, two zinc ions for HDACs 3-5 and 7-9) with and without palmitate were performed using AlphaFold 3 [45]. SCFAs and mocetinostat were docked into AlphaFold 3-generated HDAC1 models using the Swissdock attracting cavities docking engine [46–48]. Medium sampling exhaustivity and buried cavity prioritization parameters were used for all docking jobs. Random initial conditions (RIC) were set to 3 for docking SCFAs with HDAC1. RIC were set to 1 for docking mocetinostat into HDAC1 and pentanoate into HDAC4. Models were visualised and analysed using UCSF ChimeraX [49]. Palmitate molecules in the AlphaFold 3 models were truncated to C5 using UCSF ChimeraX and used for free energy calculations alongside models with palmitate. Free energy calculations were performed using PRODIGY-LIGAND [50, 51].

### HDAC activity assays

For the impact of SCFAs on specific HDAC isoforms, the fluorogenic assay for each HDAC enzyme (HDAC1-3, HDAC5) was used (BPS Bioscience). Assays were conducted according to the manufacturer’s instructions, with measurements taken in triplicates using the FLUOstar Omega plate reader.

### Metabolomic profiling

To analyze polar intracellular metabolites, CD8^+^ and CD4^+^ CAR T cells were generated as described above. After removal from the antibody on day 3, cells were put into medium containing 1 g/l ^13^C- glucose/ ^13^C- glutamine for 24 hours. Per condition, 1×10^6^ T cells were washed with 0.9 % Natriumchloride. Metabolite extraction was conducted by incubation with ice-cold 80 % methanol including internal standards. Following an incubation time of 20 min at 4 °C, cells were pelleted and the supernatant containing the polar metabolites was transferred into a new tube and stored at −80 °C until further processing. For the tracking of pentanoate, 2 mM of ^13^C-labeled pentanoate was added at day 0 following activation and samples were taken from day 1 until day 4 after activation and isolated for metabolites as described above.

Metabolites were automatically derivatized using a Gerstel MPS. Derivatization was done with 15 µl of 2 % (w/v) methoxyamine hyprochloride (Thermo Scientific) in pyridine and 15 μl N-tertbutyldimethylsilyl-N-methyltrifluoroacetamide with 1 % tert-butyldimethylchlorosilane (Regis Technologies). Measurement was carried out by GC/MS with a 30 m DB-35MS + 5 m Duraguard capillary column (0.25 mm inner diameter, 0.25 µm film thickness) equipped in an Agilent 7890B gas chromatograph (GC) connected to an Agilent 5977A mass spectrometer (MS). The GC oven temperature was held at 80□°C for 6□min and steadily adjusted at 6□°C per min until reaching 280□°C where the temperature was held for 10□min. The quadropole was set to 150 °C. The MS source operated under electron impact ionization mode at 70 eV and was held at 230°C. Targeted single ion chromatogram measurements were conducted for pyruvate (174, 175, 176, 177, 178, 179; 10 scans per second), lactate (261, 262, 263, 264, 265, 266, 267; 10 scans per second), citrate (591, 592, 593, 594, 595, 596, 597, 598, 599, 600; 10 scans per second), α-ketoglutarate (346, 347, 348, 349, 350, 351, 352, 353, 354; 10 scans per second), fumarate (287, 288, 289, 290, 291, 292, 293; 10 scans per second), glutamate (432, 433, 434, 435, 436, 437, 438, 439, 440; 10 scans per second) and malate (419, 420, 421, 422, 423, 424, 425, 426; 10 scans per second).

All chromatograms were subsequently analyzed with the MetaboliteDetector software [52].

### Preparation of cells for RNA-single cell sequencing

Cells of draining lymph nodes (dLN) and tumor-infiltrating lymphocytes (TILS) were isolated from tumor-bearing mice at day 7 and 14 post T cell infusion. Following incubation with TruStain FcX™ PLUS (anti-mouse CD16/32) antibody (Biolegend) to prevent unspecific binding, cells were stained with 2 µl of TotalSeqTM-C hashing antibodies (Biolegend) in wash buffer for 30 min at 4 °C.

Afterwards, 6 samples were pooled in one and suspended in 65 µl of PBS/0.04 % BSA.

### Single cell RNA-sequencing

Chromium™ X/iX Controller was used for partitioning single cells into nanoliter-scale Gel Bead-In-EMulsions (GEMs) and Chromium GEM-X Single Cell 5’ Kit v3 kits for reverse transcription, cDNA amplification and library construction for all gene expression, TCR and hashtag libraries (10x Genomics), following manufacturer’s instructions. A SimpliAmp Thermal Cycler was used for amplification and incubation steps (Applied Biosystems). Libraries were quantified by a QubitTM 3.0 fluorometer (Thermo Fisher Scientific) and quality was checked using a 2100 Bioanalyser with High Sensitivity DNA kit (Agilent). Libraries were pooled and sequenced using the NextSeq 2000 platform (Illumina) in paired-end mode for gene expression as well as for the T-cell receptor repertoire and hashtags. Demultiplexed FASTQ files were generated with bcl-convert v4.0.3 (Illumina). Data were analyzed using the Cell Ranger 7.2.0 software suite pipelines available on the 10xGenomics website.

The Cell Ranger output of each pool (P1 to P9) was converted to a Seurat object using the Seurat package (Read10X and CreateSeuratObject). After adding the HTO data as an independent assay, cells were demultiplexed using the HTODemux function with kmeans clustering and positive.quantile =0.99. Only singlets were selected for further analysis. The percentage of mitochondrial genes (MT-) was determined for each cell using the PercentageFeatureSet function. We filtered out low quality cells based on the number of features (< 200 or > 6000) and the percentage of mitochondrial genes (> 20 %) using the Subset function. The 9 Seurat objects were merged into a single object regrouping a total of 16908cells. Normalization and integration were performed using NormalizeData and STACAS. RunPCA, runUMAP, FindNeighbors and FindClusters with 30dimensions were used to identify 21clusters of T cells. T cells were then annotated using scGate with the pre-defined gating mouse model, and only pure T cells were subset and used for downstream analysis. A total of 4774 pure T cells were annotated for the different CD4^+^ and CD8^+^ T cell subsets using ProjecTILs and the human reference atlas to characterize cell states.

### *In vivo* models

For all experiments, group sizes were determined based on experience by previously published models. Tumor-bearing mice were randomly assigned to receive CAR T cell infusions, to ensure similar tumor sizes across groups before treatment. Tumor engraftment and T cell infusions were performed by blinded technicians. Generated CAR T cells were not sorted but additional non-transduced cells were added, so that the same amount of total T cell count was applied to each animal.

### MM5080 tumor models

For adoptive transfer experiments with MM5080 tumor cells, mice were intravenously (i.v.) injected with 5×10^6^ tumor cells. On day 10 after tumor injection, 0.5×10^6^ CD8^+^ and CD4^+^ CAR each T cells were transferred i.v into Bl6 mice irradiated with 2.5 Gy TBI. IL-2 was injected for 4 consecutive days (20.000U/mouse).

### Panc02 tumor models

For adoptive transfer experiments with Panc02 mROR1 tumor cells, mice were subcutaneously (s.c.) injected with 2×10^6^ tumor cells. On day 5 after tumor injection, 2×10^6^ CD8^+^ CAR T cells were transferred i.v into Bl6 mice.

For the analysis of transferred CAR T cells by scRNA sequencing, mice were subcutaneously (s.c.) injected with 1×10^6^ tumor cells. After an inoculation time of 7 days, 1×10^6^ CAR T cells with a 1:1 mixture of CD4:CD8 CAR T cells were transferred into tumor bearing mice. On day 7 and day 14 after T cell injection, 5 mice per group were sacrificed and samples of dLN and TILs were sent for RNA single cells sequencing.

### MC38 tumor model

For adoptive transfer experiments with MC38 mROR1 tumor cells, mice were subcutaneously (s.c.) injected with 2×10^6^ tumor cells in PBS with matrigel. Mice were treated with 2×10^6^ CD8^+^ CAR T cells on day 7.

For all solid tumor *in vivo* experiments, tumor progression was monitored by caliper every second day. Mice were monitored daily and euthanized when mice reached limits designated in the approved protocols including maximum tumor burden. At the endpoint of the experiments, peripheral blood was obtained by tail vein puncture. Cell pellets were resuspended in ACK lysis (Thermo Fisher) buffer for 10 min. Spleen, bone marrow, draining lymph node and tumors were collected. After determination of the weight, tumors were dissociated using the tumor dissociation kit (Miltenyi Biotech) and a GentleMACSTM Dissociator (Miltenyi Biotech) according to manufacturer’s constructions. All organs were passed through a 70 µM strainer to acquire single-cell suspensions. Tumor-infiltrating lymphocytes were isolated using CD45 (TIL) microbeads (Miltenyi Biotech). Consequent cell suspensions were washed with PBS and stained with live/dead marker (Biolegend) for 20 min at RT. After washing, samples were incubated with TruStain FcX™ (anti-mouse CD16/32) Antibody (Biolegend) to prevent unspecific binding for 15min at RT and then stained with FACS antibodies for analysis (List of antibodies in **Suppl. Table 3**).

### Quantification and statistical analysis

The results are shown as mean ± standard error of the means (SEM). To determine the statistical significance of the differences between two experimental groups unpaired Student’s t tests were performed using Prism 9 software (GraphPad). For identify the statistical significance of the difference between more than to experimental groups one-way or two-way analysis of variance (ANOVA) with Tukey’s multiple-comparison test were performed using Prism 9 software (GraphPad). For *in vivo* tumor growth curves, significance was determined at indicated timepoints on the blot by unpaired Student’s t-test comparing control group with the treatment group. Significance for the survival data was calculated using the log-rank Mantel-Cox test. Samples sizes were based on experience and complexcity of the experiment but no methods were used to determine normal distribution of the samples. Differences reached significance with p- values ≤0.05, p ≤ 0.0.1, p ≤ 0.001 or ≤ 0.0001. The figure legends contain the number of independent experiments or mice per group that were used in the respective experiment.

For the clinical study, statistical analyses were conducted using RStudio version 2023.12.1+402 (Posit, Boston, MA, USA) and R version 4.3.3 (The R foundation, Vienna, Austria). The level of significance was set at a two-sided p ≤ 0.05 with 95% confidence intervals. Grouped data are presented as violin plots. To compare two groups, based on the distribution of the data, the Wilcoxon-Mann-Whitney-Test (for non-normally distributed data) or the t-test (for normally distributed data) were conducted and adjusted for false discovery rate (FDR) using the rstatix package (version 0.7.2). Progression-free survival (PFS) was defined as the time from CAR T cell infusion to disease progression or death, whichever occurred first. Overall survival (OS) is defined as the timespan between the CAR T cell infusion and the patient’s death irrespective of the cause. Metabolite cutoffs were determined utilizing surv_cutpoint () function from the survminer package (version 0.4.9). Survival data are depicted as Kaplan-Meier curves. Differences between two groups were assessed with the log-rank test via the ggsurvfit () function and package (version 1.0.0). For comparisons of more than one group, a cox regression was performed using the coxph() function from the survival package (version 3.5-8).

## Data availability

The authors declare that data supporting the findings of this study are available within the paper and its supplementary information files. Data derived from single cell sequencing experiments will be made publicly available upon manuscript acceptance.

## Acknowledgments

The study was supported by the Austrian Science Fund project SFB F70 and by the Horizon 2020 Marie Slodowska Curie Innovative Training Network “Enlight-ten+”(Grant 955321 to WE), the Instituto de Salud Carlos III (PI22/00983 to JAMC) co-financed by European Regional Development Fund-FEDER “A way to make Europe” Red de Terapias Avanzadas TERAV (RD21/0017/0009 to FP, PMPTA22/00109 to JRR-M), Ministerio de Ciencia e Innovación co-financed by European Regional Development Fund-FEDER “A way to make Europe” (PID2022-137914OB-I00 to JRR-M, PID2021-128283OA-I00 to TL, PID2022-137265OB-I00 to JJL), Centro de Investigación Biomédica en Red de Cáncer CIBERONC (CB16/12/00489 and CB16/12/00369 to FP; CB16/12/00489, and CB16/12/00369 to JAMC), European Commission (H2020-JTI-IMI2-2019-18: Contract 945393; SC1-PM-08-2017: Contract 754658 to FP), the Engineered Air Chair in Cancer Research from University of Calgary (to PN), the European Molecular Biology Conference (EMBC) under EMBO Installation (Grant agreement number 5342-2023 to PP), the Ministry of Education, Science and Sports of Lithuania (Measure No. 12-001-01-01-01, Project No. S-A-UEI-23-10 to PP), the German Jose Carreras Foundation, the National Institutes of Health/National Cancer Institute (NIH/NCI) Memorial Sloan Kettering Cancer Center Support Grant (P30 CA008748 to RS), NIH-NCI K08-CA282987 (to RS) and the Research Council of Lithuania (LMTLT) (grant agreement number S-PD-24-74 to AS).

Moreover, support was received from the Innovative Medicines Initiative 2 Joint Undertaking, from the European Union’s Horizon 2020 research and innovation program, EFPIA and the European Hematology Association (EHA) (grant agreement No. 116026, T2EVOLVE to LW, DB, FP, MH and ML), the Wilhelm-Sander-Stiftung (Grant No. 2022.134.1 to AV, KM and ML), ERA-NET TRANSCAN-3 (EC co-funded call 2021, SmartCAR-T to KZM, PN, MH and ML), the Paula & Rodger Riney Foundation (to MvdB, MH, JAMC and ML), IZKF Würzburg (S-511 to SS), the Fundacion Arnal Planelles (to JAMC) the German Research Foundation (Deutsche Forschungsgemeinschaft, DFG, SFB-1454 (project 432325352 to KH), SFB-1371 (Project 395357507 to HP and KK), TRR 221 (subproject A03 to MH, HE and ML; subproject A08 to HP and MP; and TRR338, (subproject A02; to MH, HE and ML, seed grant; to ML), the German Cancer Aid (70114547 to HP), the Bavarian Cancer Research Center (BZKF) (to MP, MAF, MH, ML and HP) and the ReforM program of the University Regensburg (to MP). This work was co-funded by the European Union (project MICROBOTS, Grant No. 101124680 to HP). We thank the Core Unit SysMed at the University of Würzburg for excellent technical support, scRNA-seq data generation and analysis. This work was supported by the IZKF at the University of Würzburg (project Z-6). We also thank all BayBioMS members for their help and fruitful discussions. The TripleTOF 6600 (Sciex) mass spectrometer and the Q Exactive HFX (Thermo) was funded in part by the German Research Foundation (INST 95/1434-1 FUGG and INST 95/1435-1 FUGG).

## Author Contributions

Study design: ML, MH, MVDB and AV; Core *in vitro* experiments: SS, JF, FN; Supporting *in vitro* experiments NAG, KZM, PB, FF, ZR, PHT, PSMU, JRRM, *in vivo* studies: SS, JF, LW, TL, MLA; data analysis and modeling: SS, FN, TF, MB, KH, NL, XKY, PN, AS and PP; clinical study design and conduction: JS, CST, MS, KK, MP, MAF, HP; writing - original draft: ML, SS; writing – review and editing: RS, JJS, JAMC, FP, WE, HE, PN MH, MP, HP, KM, PP; funding acquisition: FP, ML, MH, HP; supervision, FP, ML, MH, HE, FP, WE, JW, DB, PP, AV, KH, PN, MVDB, HP, JJL, JAMC

