## Extended Data Figures for "Metabolization of microbial postbiotic pentanoate drives anti-cancer CAR T cells"

Staudt et al.

**Extended Data Figures**

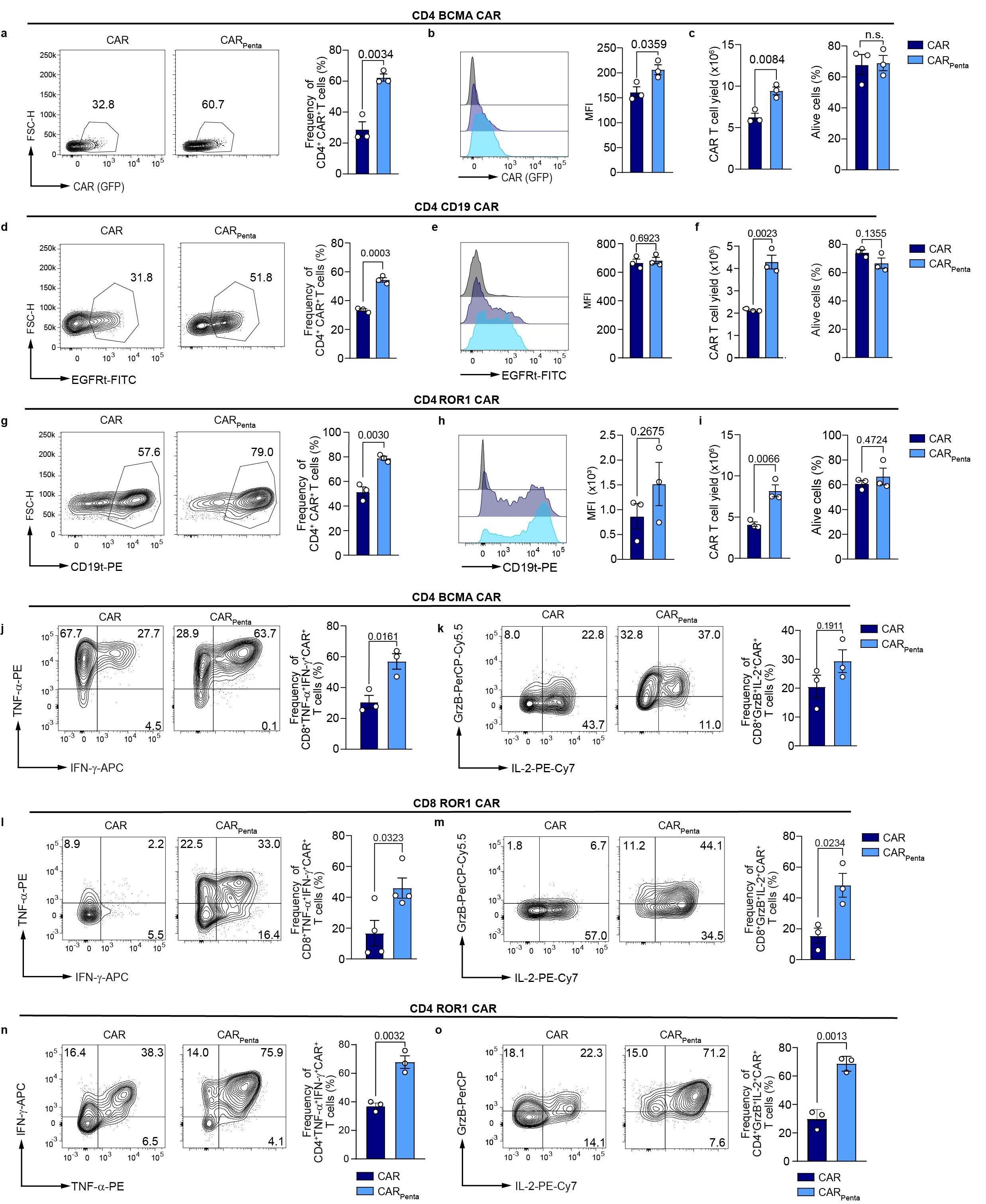

**Extended Data Figure 1: Pentanoate treatment enhances generation of multiple CAR constructs. a**, Dot blot and bar graph show expression of transduction marker of CD4^+^ BCMA - CAR T cells on day 6. Mean ± SEM from *n = 3*. **b**, Representative histogram and mean fluorescence intensity (MFI) of the transduction marker. Mean ± SEM from *n = 3*. **c**, CAR T cell yield and live cells of BCMA^+^ - CAR T cells on day 7. Mean ± SEM from *n = 3*. **d**, Dot blots and bar graph show tEGFR expression of CD4^+^ CD19- CAR T cells on day 6. Mean ± SEM from *n = 3*. **e**, MFI for generated CAR T cells as histogram representative and bar graph. Mean ± SEM from *n = 3*. **f**, Cell number and viability of cells of generated CAR^+^ T cells on day 7. Mean ± SEM from *n = 3*. **g**, Dot blots and bar graph show expression of CD4^+^ ROR1- CAR T cells on day 6. Mean ± SEM from *n = 3*. **h**, MFI of the transduction marker for generated CAR T cells as histogram representative and bar graph. Mean ± SEM from *n = 3*. **i**, CAR^+^ T cell number and viability on day 7. Mean ± SEM from *n = 3*. **j** and **k**, Representative dot plot and bar graph show frequency of IFN-γ^+^TNF-α^+^ double positive (**i**) and GranzymeB^+^IL-2^+^ double positive (**j**) CD4^+^ BCMA- CAR T cells upon restimulation. Mean ± SEM from *n = 3*. **l** and **m**, Dot plot and bar graph show frequency of IFN-γ^+^TNF-α^+^ double positive (**l**) and GranzymeB^+^IL-2^+^ double positive (**m**) upon restimulation of CD8^+^ ROR1- CAR T cells. Mean ± SEM from *n = 3*. **n** and **o**, CD4^+^ ROR1 – CAR T cells were restimulated for 5h with brefeldin A. Representative dot plot and bar graph show frequency of IFN-γ^+^TNF-α^+^ double positive (**n**) and GranzymeB^+^IL-2^+^ double positive (**o**) cells. Mean ± SEM from *n = 3*.

(**a**-**h**) Data represent pooled data from 3 independent experiments. Statistical analysis was performed using unpaired two-tailed Student’s *t* test.

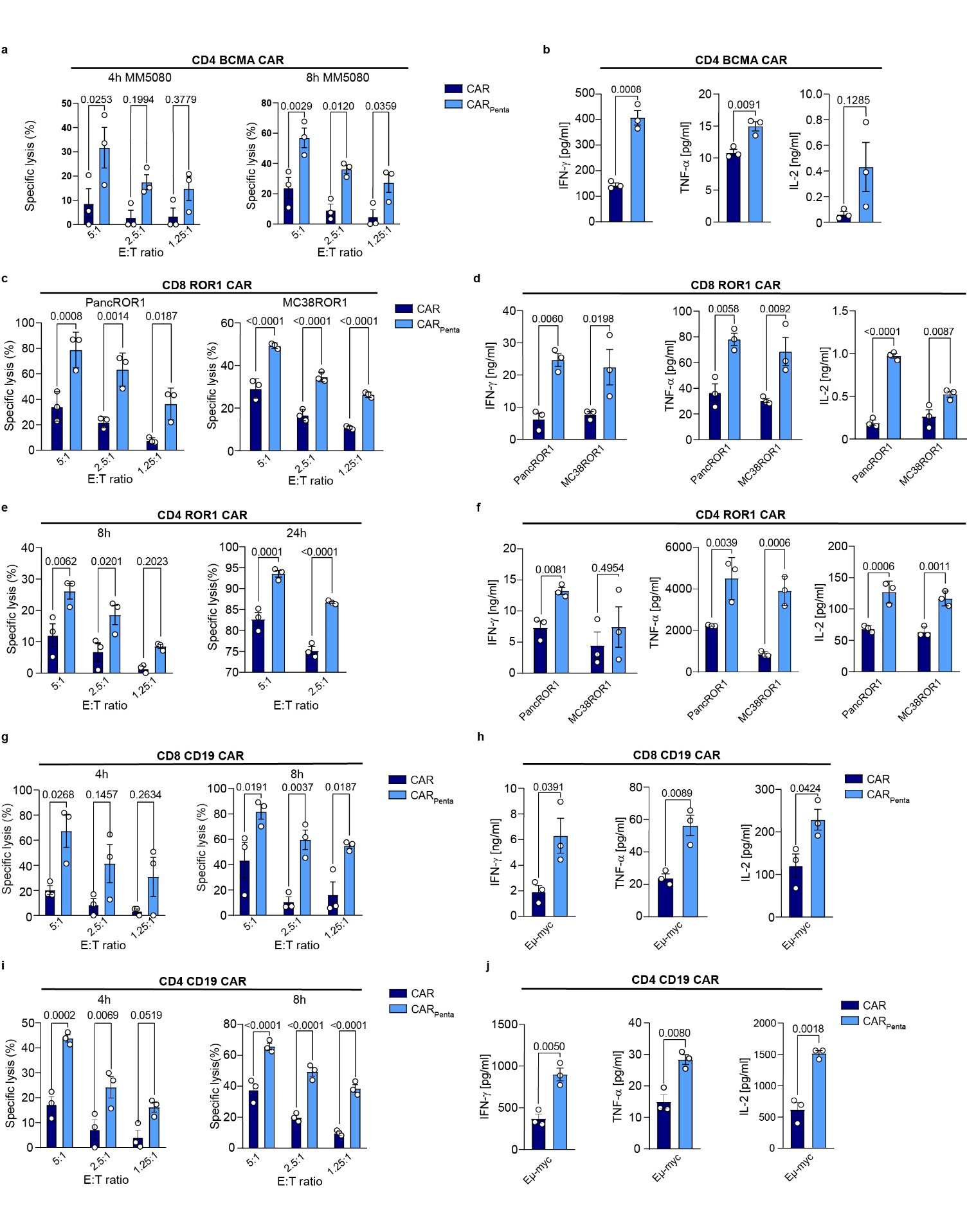

**Extended Data Figure 2: Functional assessment of generated CAR T cells. a**, Specific cytolytic activity of CD4^+^ BCMA – CAR T cells against MM5080 tumor cells after 4 and 8h. Mean ± SEM from *n = 3*. b, Secretion of IFN-γ, TNF-α and IL-2 of CD4^+^ BCMA – CAR T cells after co-incubation with target cells for 24h determined by ELISA. Mean ± SEM from *n = 3*. **c**, Specific lysis of CD8^+^ ROR1 – CAR T cells of PancROR1 and MC38ROR1 target cell lines after h. Mean ± SEM from *n = 3*. **d**, Cytokine secretion of CD8^+^ ROR1 – CAR T cells was measured after co-incubation with target cell lines for IFN-γ, TNF-α and IL-2 after 24h by ELISA. Mean ± SEM from *n = 3*. **e**, Cytolytic activity of CD4^+^ ROR1 – CAR T cells against antigen-expressing cell line after 8h and 24h. Mean ± SEM from *n = 3*. **f**, IFN-γ, TNF-α and IL-2 concentration in the supernatant was measured by ELISA following the co-incubation of CD4^+^ ROR1 – CAR T cells with antigen-expressing tumor cells lines for 24 h. Mean ± SEM from *n = 3*. **g**, Specific lysis of CD19-expressing Eµ-myc tumor cell lines by CD8^+^ CD19 – CAR T cells after 4 and 8h was determined. Mean ± SEM from *n = 3*. **h**, Cytokine secretion of IFN-γ, TNF-α and IL-2 was detected by ELISA following 24-hour co-incubation with CD8+ CD19 – CAR T cells and antigen-expressing tumor cells. Mean ± SEM from *n = 3*. **i**, Specific lytic activity of CD4+ CD19 – CAR T cells against Eµ-myc tumor cells after 4 and 8h. Mean ± SEM from *n = 3*. **j**, Cytokine secretion of IFN-γ, TNF-α and IL-2 of CD4^+^ CD19 – CAR T cells was determined by ELISA. Mean ± SEM from *n = 3*.

(**a** - **j**) Data show pooled data from 3 independent experiments with *n=3* biological replicates; pooled data from *n=3* independent experiments; mean ± SEM was calculated for *n=3* independent experiments. Statistical analysis was performed using two-way analysis of variance (ANOVA) with Tukey’s multiple-comparison test.

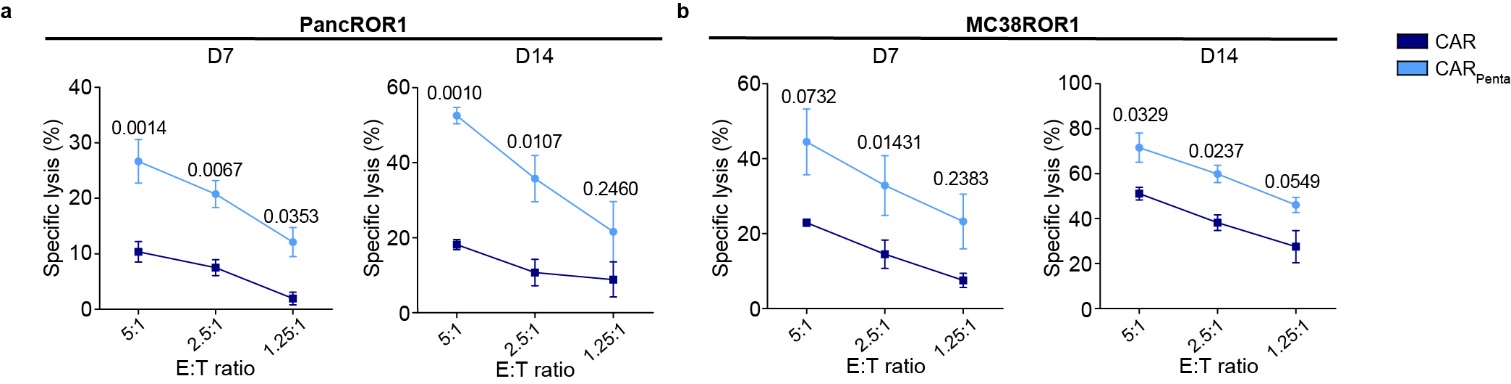

**Extended Data Figure 3: ROR1 CAR T cell killing day 14. a** and **b,** Specific lysis of generated CD8^+^ ROR1- CAR T cells against PancROR1 (**a**) and MC38ROR1 (**b**) target cells on day 7 or day 14 after treatment. Mean ± SEM from *n = 3*.

Statistical analysis was performed using one-way analysis of variance (ANOVA) with Tukey’s multiple-comparison test (**b** and **c**).

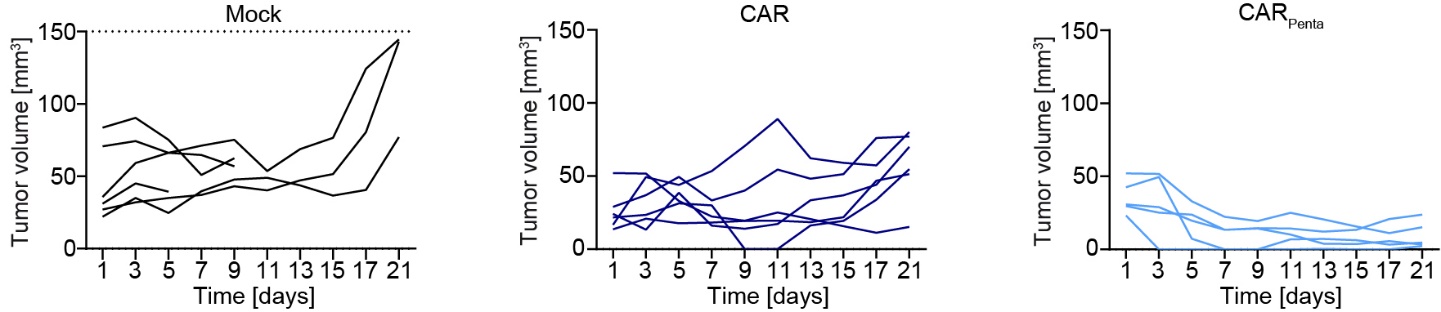

**Extended Data Figure 4: Organ analysis ex vivo.** MC38ROR1 tumor volume for mock, CAR T cells or CAR_Penta_ T cells over the experimental course.

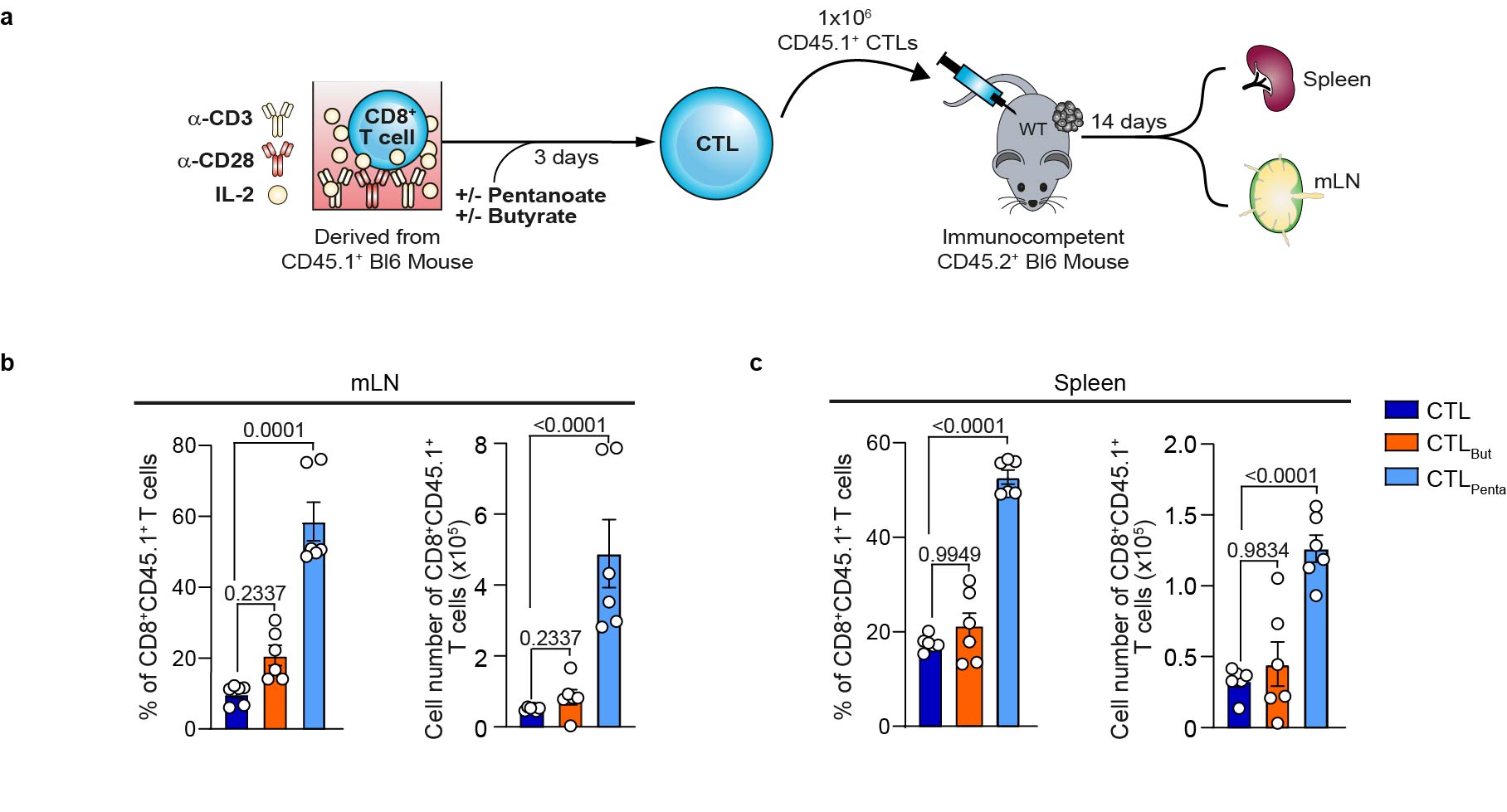

**Extended Data Figure 5: Polyclonal in-vivo expansion. a**, Experimental setup of in-vivo experiment with polyclonal T cells with stated treatments. **b** and **c,** Percentage and total cell number of CD8^+^ T cells in the lymph node (**b**) and spleen (**c**). Mean ± SEM from *n = 6* mice/group.

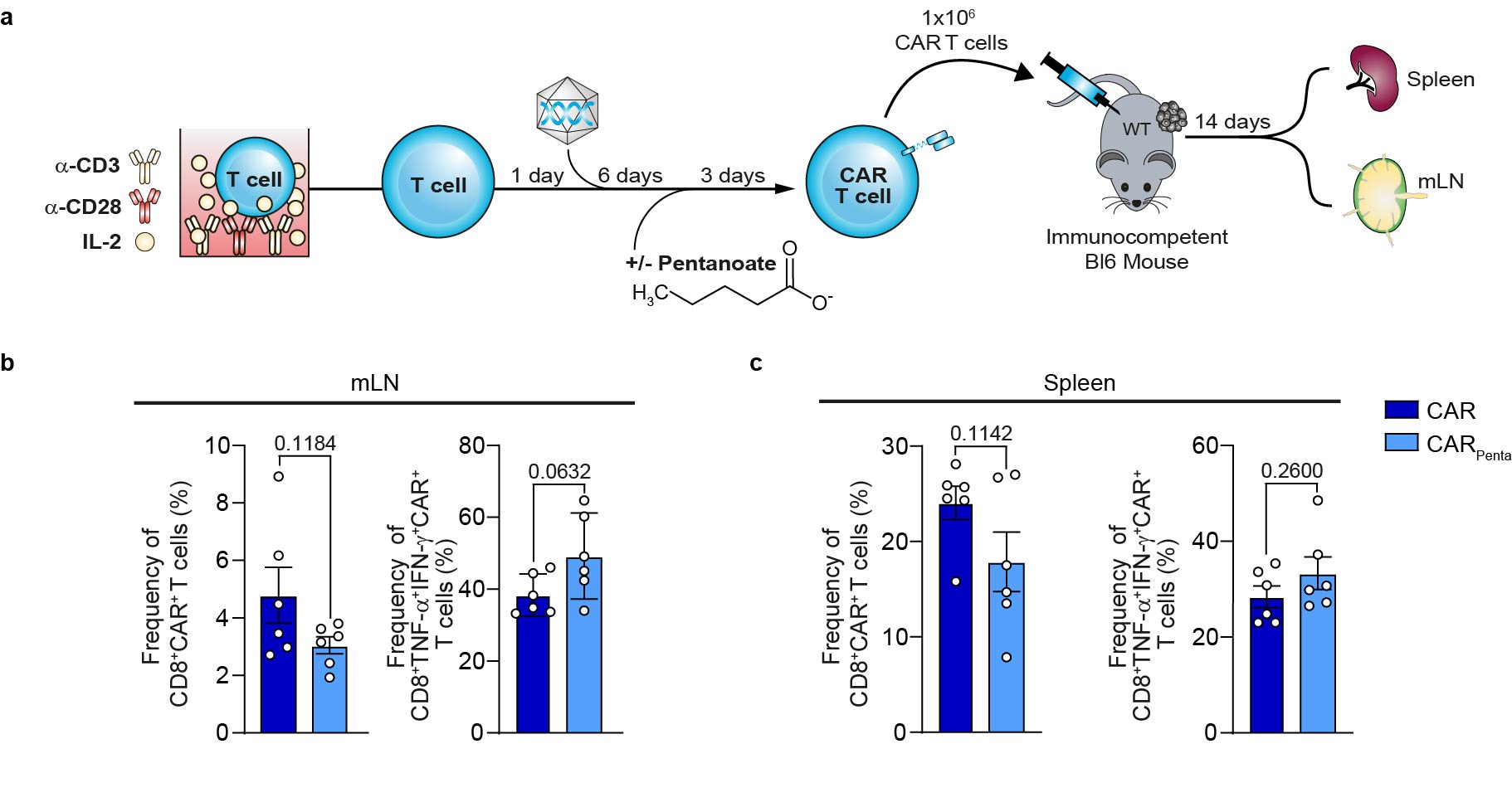

**Extended Data Figure 6: In vivo expansion of CAR T cells after late treatment. a**, Experimental design for the analysis of *in vivo* expansion of CAR T cells following Pentanoate treatment after CAR T cell generation. **b** and **c**, Flow cytometry staining identify the percentage of ROR1- CAR T cells and cytokine secretion of TNF-α and IFN-γ in the lymph node (**b**) and spleen (**c**) following restimulation at the endpoint of the experiment; Mean ± SEM from *n* = 6 mice/group.

Statistical analysis was performed using unpaired two-tailed Student’s *t* test (**b** and **c**).

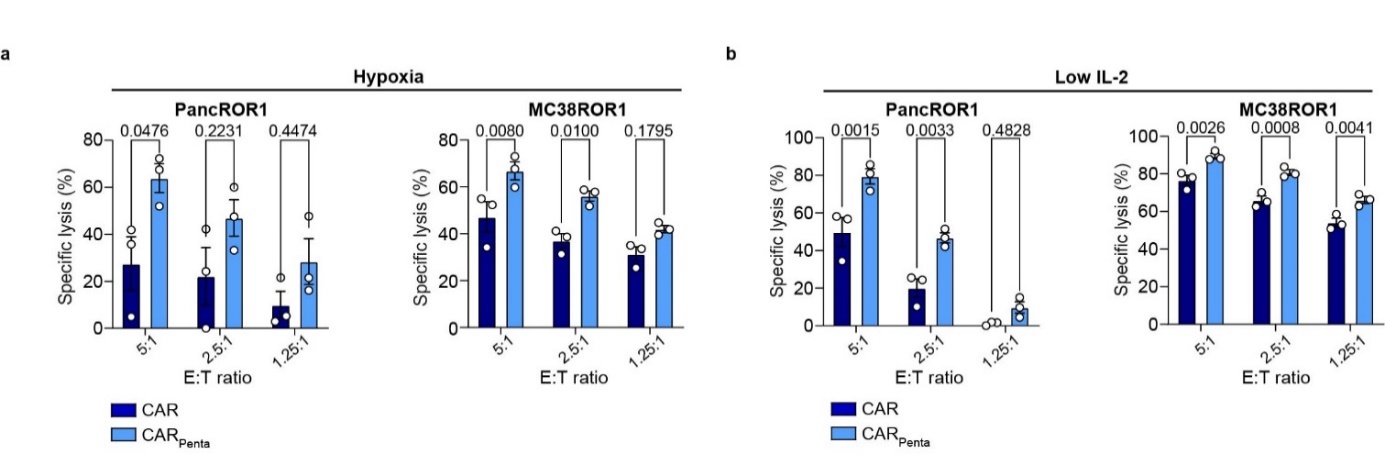

**Extended Data Figure 7: Hypoxia and low IL-2. a**, Cytolytic activity of generated CD8^+^ CAR T cells against PancROR1 and MC38ROR1 tumor cells at variable E:T ratios after 6 h under hypoxia conditions. **b**, Specific lysis of antigen-presenting tumor cells by CD8^+^ CAR T cells generated with low IL-2 conditions after 24 h. Mean ± SEM from *n = 3*.

Statistical analysis was performed using one-way analysis of variance (ANOVA) with Tukey’s multiple-comparison test (**b** and **c**).

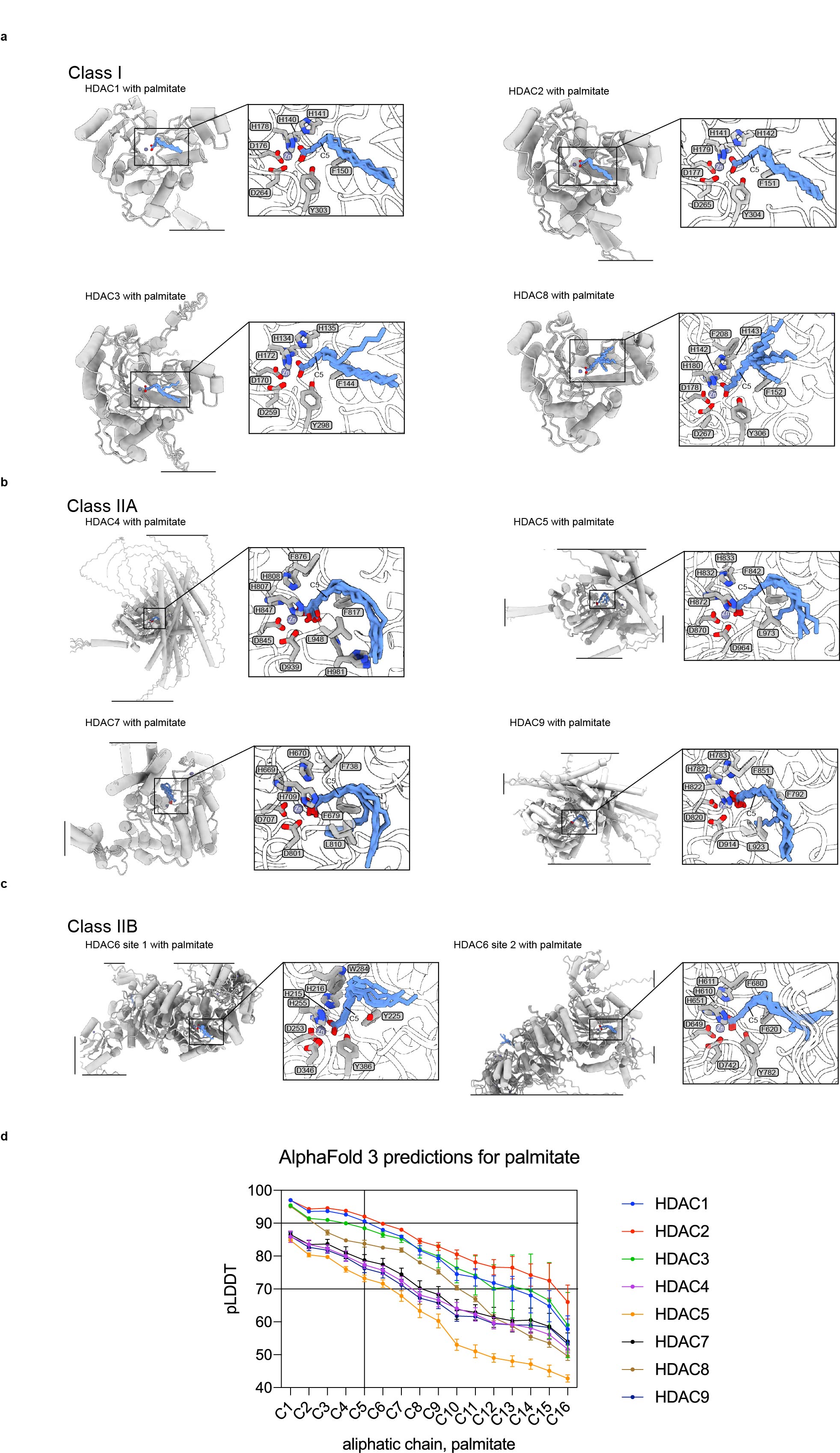

**Extended Data Figure 8: Prediction of HDAC models bound to zinc and palmitate.** Shown are superimpositions of AlphaFold 3-predicted structures of HDACs (gray cartoons) bound to zinc (spheres) and palmitate (blue sticks). Five predicted structures were superimposed for each HDAC of classes I (**a**), IIA (**b**) and IIB (**a**). **d**, Shown are per-carbon-atom local confidence pLDDT values for the aliphatic chain of palmitate, modeled into HDACs of classes I (HDACs 1, 2, 3 and 8) and IIa (HDACs 4, 5, 7 and 9). Mean ± s.d. of n = 5 AlphaFold 3 predictions.

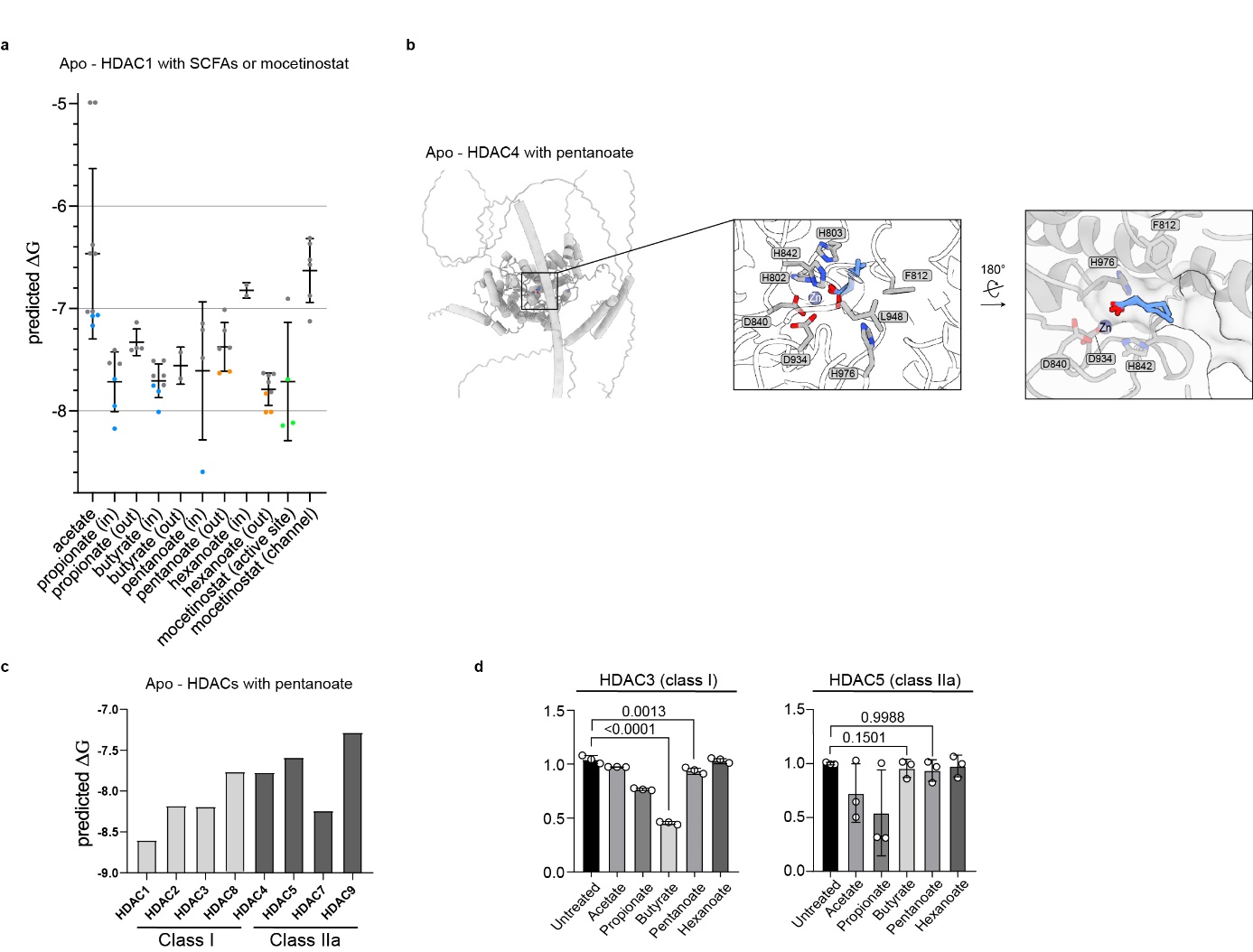

**Extended Data Figure 9: Swissdock SCFAs. a**, Free energy ΔG predictions of SCFAs and mocetinostat docked into an AlphaFold 3-predicted model of zinc bound HDAC1. Ten lowest free energy ΔG conformations were plotted separately based on the conformation relative to the catalytic center (in and out; active site and channel); Mean ± s.d.. **b**, Pentanoate was docked into AlphaFold 3-predicted models of zinc-bound HDACs. **c**, Predicted binding energies for pentanoate into HDAC 1-5 and 7-9 proteins. ΔG values for the best fitting orientation are shown. Proteinmodel was obtained using AlphaFold 3. Docking was performed using SwissDock. **d**, Inhibition of HDAC 3 and HDAC 5 for several SCFAs. Mean ± SEM from *n = 3*.

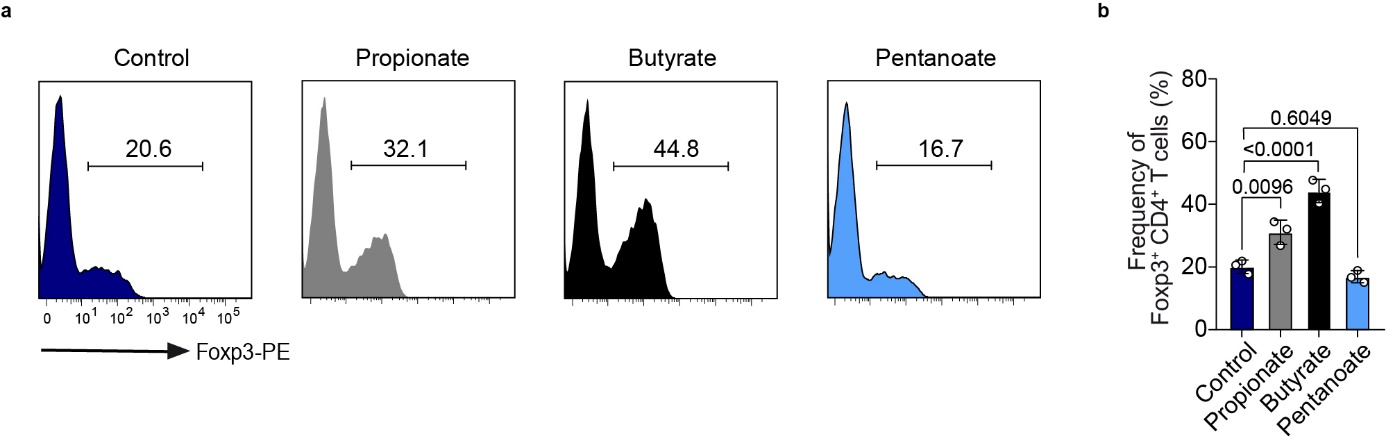
**Extended Data Figure 10: Induction of regulatory T cell phenotype by SCFAs. a**, Staining for FoxP3 following incubation of CD4^+^ T cells with indicated SCFAs as representative histograms. **b**, Bar graph of frequency of FoxP3^+^CD4^+^ T cells. Mean ± SEM from n=3.

Statistical analysis was performed using two-way analysis of variance (ANOVA) with Tukey’s multiple-comparison test (**b**).

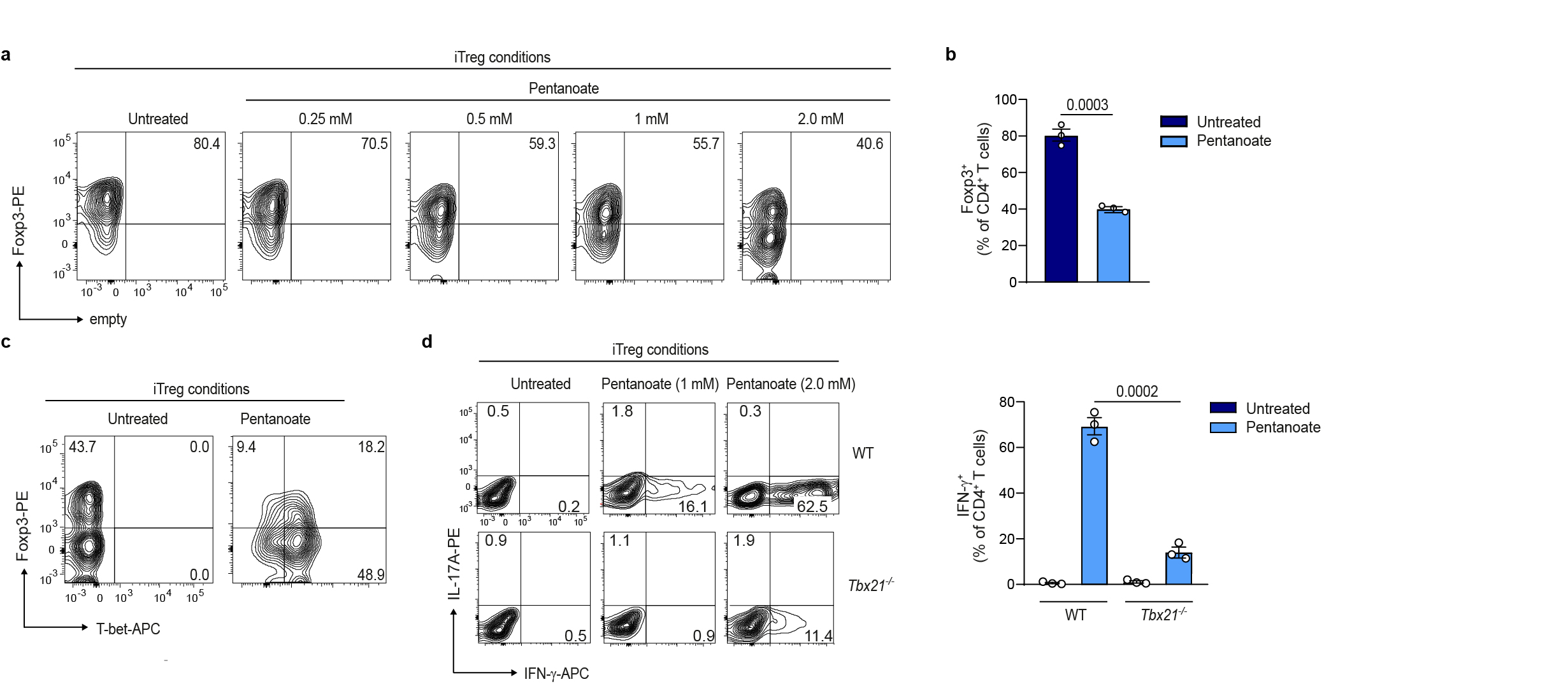

**Extended Data Figure 11: Repression of Regulatory T cell phenotype by pentanoate. a**, Representative flow cytometry blots of FoxP3 in CD4^+^ T cells following the incubation with pentanoate with several concentrations for 3 days. **b**, Bar graph for FoxP3 expression for untreated T cells or treated with 2mM pentanoate. Mean ± SEM from *n = 3*. **c**, Representative staining for FoxP3^+^T-bet^+^ of CD4^+^ T cells following iTreg induction. **d**, Dot blot representative of IL-17A^+^IFNy^+^ double positive CD4^+^ T cells generated under iTreg conditions either from wildtype of Tbx21^-/-^ mice. Graph shows IFN-y^+^ CD4^+^ T cells. Mean ± SEM from *n=3*.

Statistical analysis was performed using unpaired two-tailed Student’s *t* test (**b**) or two-way analysis of variance (ANOVA) with Tukey’s multiple-comparison test (**d**).

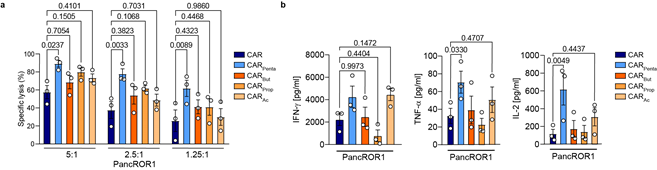

**Extended Data Figure 12: Modulation of SCFAs for CD8^+^ CAR T cell generation. a**, Cytolytic activity at various E:T ratios of ROR1- CAR T cells generated with indicated metabolites. Mean ± SEM from *n = 3*. **b**, Cytokine secretion for IFN-γ, TNF-α and IL-2 was measured for generated CAR T cells following the co-incubation with antigen-expressing tumor cells at a 5:1 E:T ratio after 24 h. Mean ± SEM from *n = 3*.

Statistical analysis was performed using two-way analysis of variance (ANOVA) with Tukey’s multiple-comparison test (**a** and **b**).

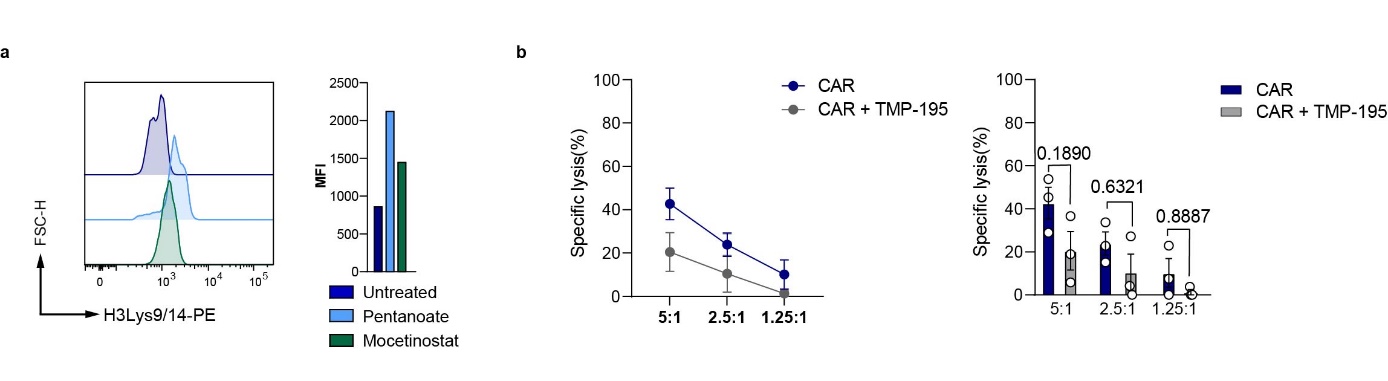
**Extended Data Figure 13: Histone acetylation of CD4^+^ CAR T cells. a**, Staining of H3 (Lys9/14) following the treatment of CD4^+^ T cells with pentanoate or mocetinostat. 1 out of 3 performed experiments is shown. **b**, Specific lysis of CD8^+^ CAR T cells either untreated or treated with TMP-195 at different E:T ratios after 6 h. Control CAR same as Figure 3. Mean ± SEM from *n = 3*.

Statistical analysis was performed using two-way analysis of variance (ANOVA) with Tukey’s multiple-comparison test (**b**).

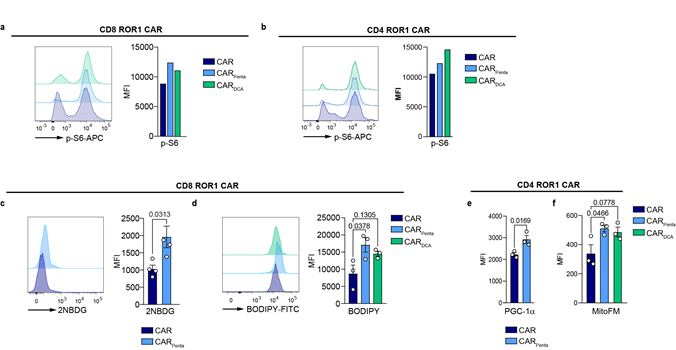

**Extended Data Figure 14: Pentanoate and DCA induce metabolic changes. a** and **b**, Following incubation with T cells in the presence of pentanoate or DCA, phospho-S6 levels were measured for CD8^+^ (**a**) and CD4^+^ (**b**) CAR T cells by flow cytometry. 1 out of 3 performed experiments is shown. **c**, Measurement of 2-NBDG in CD8^+^ CAR T cells. Mean ± SEM from *n = 3*. **d**, BODIPY-staining of CAR T cells generated with indicated metabolites. Mean ± SEM from *n = 3*. **e** and **f**, Mean fluorescence intensity (MFI) of PGC-1α (**e**) and mitochondrial mass (**f**) for CD4^+^ CAR T cells following the generation in pentanoate- or DCA- supplemented medium. Mean ± SEM from *n = 3*.

Statistical analysis was performed using unpaired two-tailed Student’s *t* test (**c** left panel,**f**) or one-way analysis of variance (ANOVA) with Tukey’s multiple-comparison test (**e** right panel).

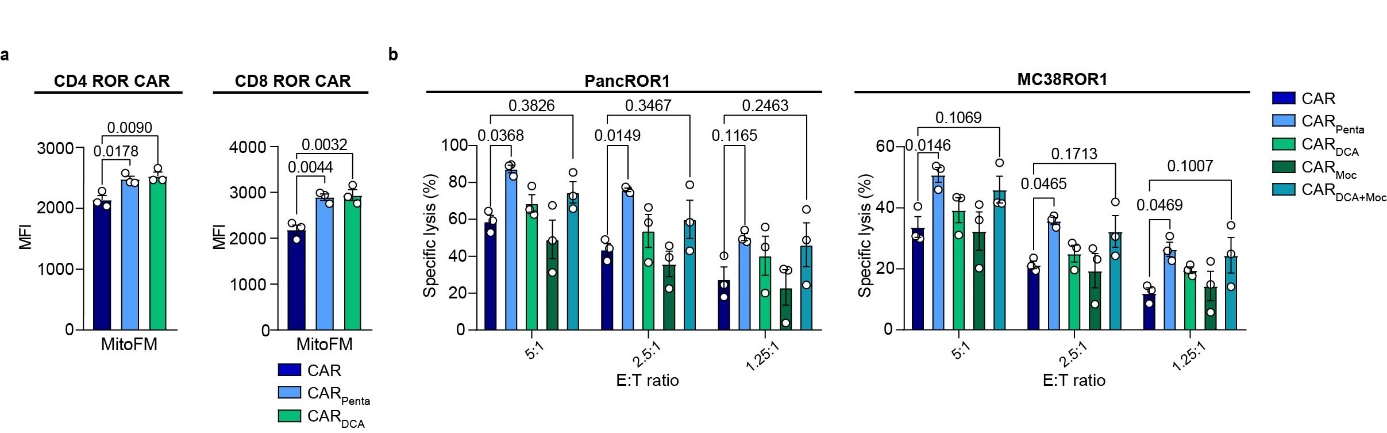
**Extended Data Figure 15: Pentanoate affects CAR T cells with a CD28 costimulatory domain. a**, MitoFM staining for ROR1- CAR T cells. Mean ± SEM from *n = 3*. **b**, Cytolytic activity of CD8^+^ CAR T cells against antigen-expressing tumor cell lines at different E:T ratios after 4 h. Mean ± SEM from *n = 3*.

Statistical analysis was performed using one-way ANOVA (**a**) or two-way analysis of variance (ANOVA) with Tukey’s multiple-comparison test (**b**).

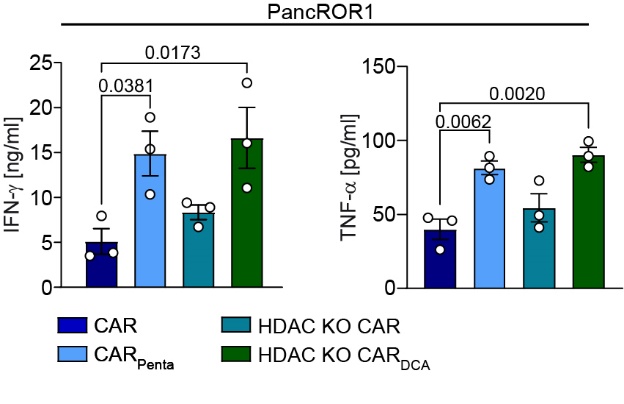

**Extended Data Figure 16: HDAC KO T cells cytokines. a**, Secretion of IFN-γ and TNF-α of generated CD8^+^ CAR T cells after co-incubation with PancROR1 tumor cells at a 5:1 E:T ratio for 24 h. Mean ± SEM from *n = 3*.

Statistical analysis was performed using one-way analysis of variance (ANOVA) with Tukey’s multiple-comparison test (**a**).

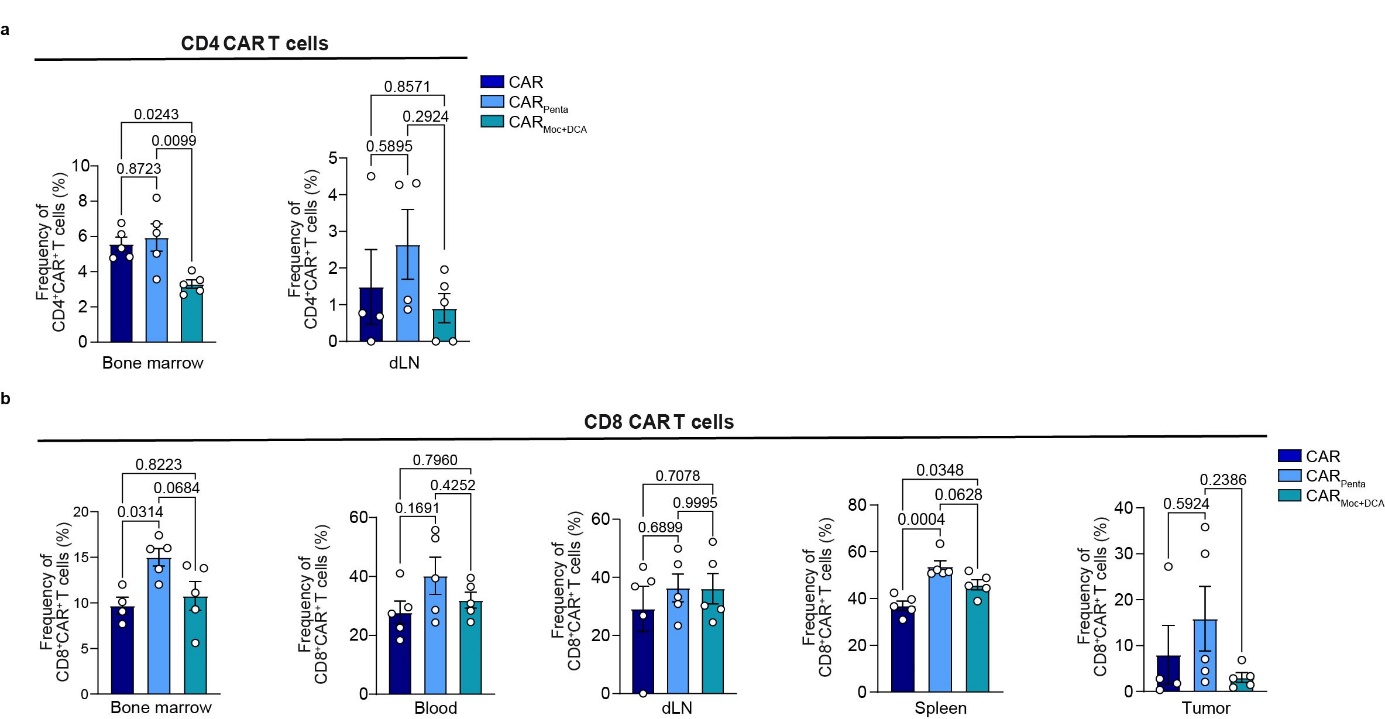

**Extended Data Figure 17: Expansion in-vivo of Pentanoate or DCA/Mocetinostat treated CAR T cells. a**, Frequency of CD4^+^ ROR1- CAR T cells in the bone marrow and draining lymph node (dLN) on day 14. **b**, Frequency of CD8^+^ ROR1- CAR T cells in the bone marrow, blood, dLN, spleen and tumor on day 14. Mean ± SEM from *n = 5* mice/group.

Statistical analysis was performed using two-way analysis of variance (ANOVA) with Tukey’s multiple-comparison test (**a** and **b**).

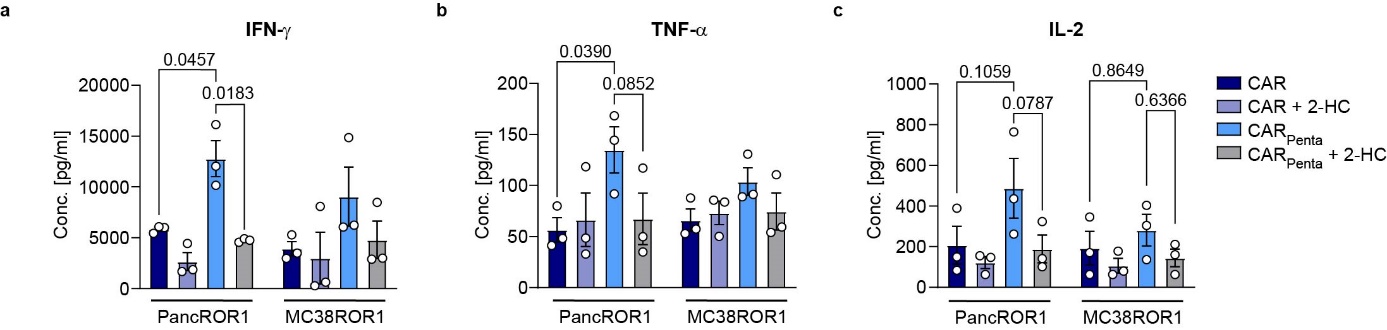

**Extended Data Figure 18: Cytokine secretion for CAR T cells treated with 2-HC.** CD8^+^ CAR T cells were co-incubated with antigen-expressing tumor cells for 24 h. Cytokine secretion for IFN-γ (**a**), TNF-α (**b**) and IL-2 (**c**) was determined by ELISA. Mean ± SEM from *n = 3*.

Statistical analysis was performed using two-way analysis of variance (ANOVA) with Tukey’s multiple-comparison test (**a**-**c**).

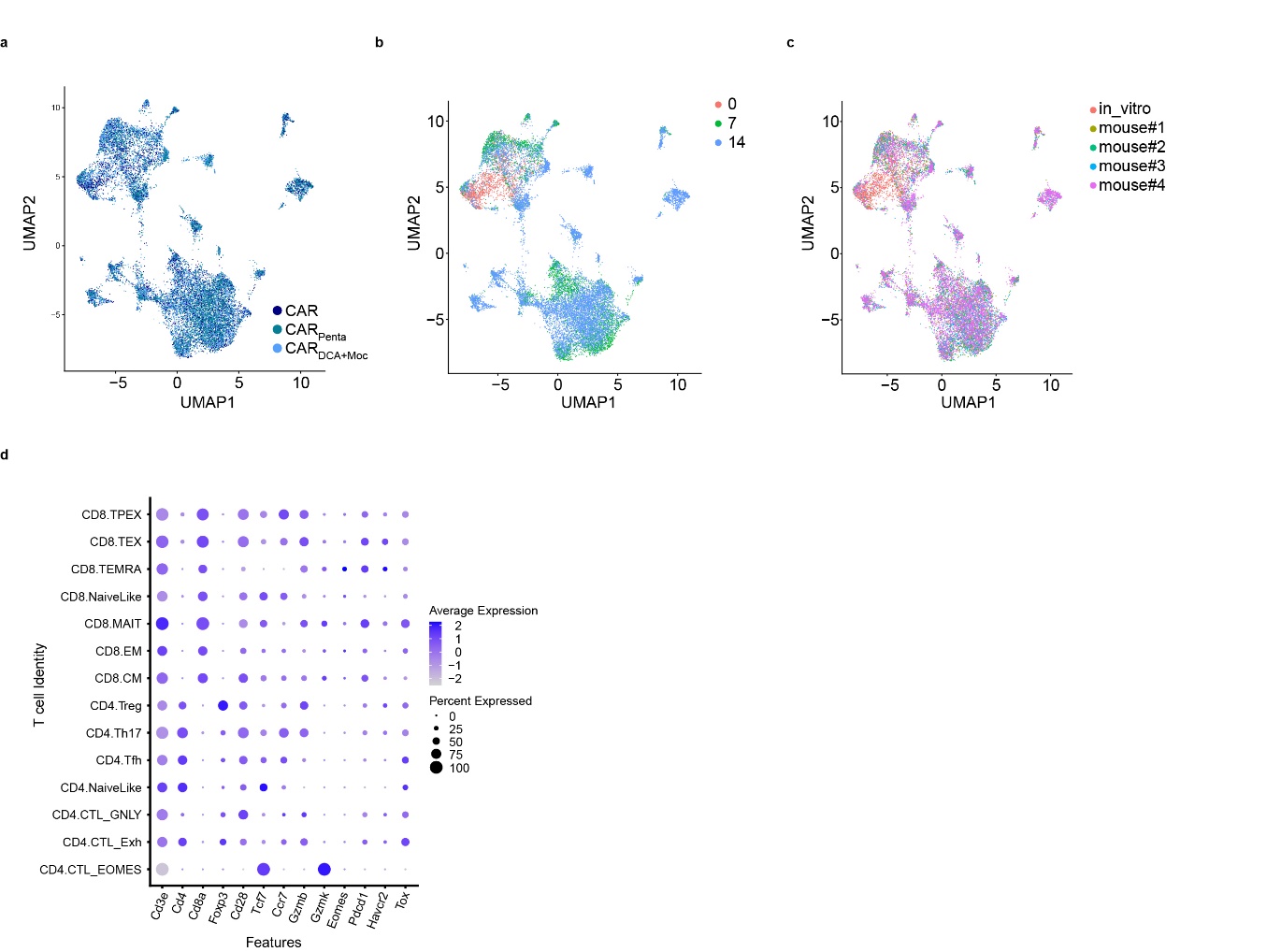

**Extended Figure 19: Singe cell sequencing data ex vivo. a** - **c**, Uniform Manifold Approximation and Projection (UMAP) showing the distribution of immune cells collected from mice using scRNAseq data in the LSI space. Each point represents one cell. The cells are marked by color code based on conditions (**a**), mouse of origin (**b**) and day/time points (**c**). **d**, Dot plot representing the scRNAseq expression of key genes involved in T cell biology and differentiation (CD3, CD4, CD8, Foxp3, CD28, Tcf7, CCR7, Gzmb, Gzmbk, Eomes, Pdcd1, Havcr2, Tox) measured in each subset of T cells. The dot size represents the percentage of cells with values detected in each subset. The color represents the average gene expression in each subset.

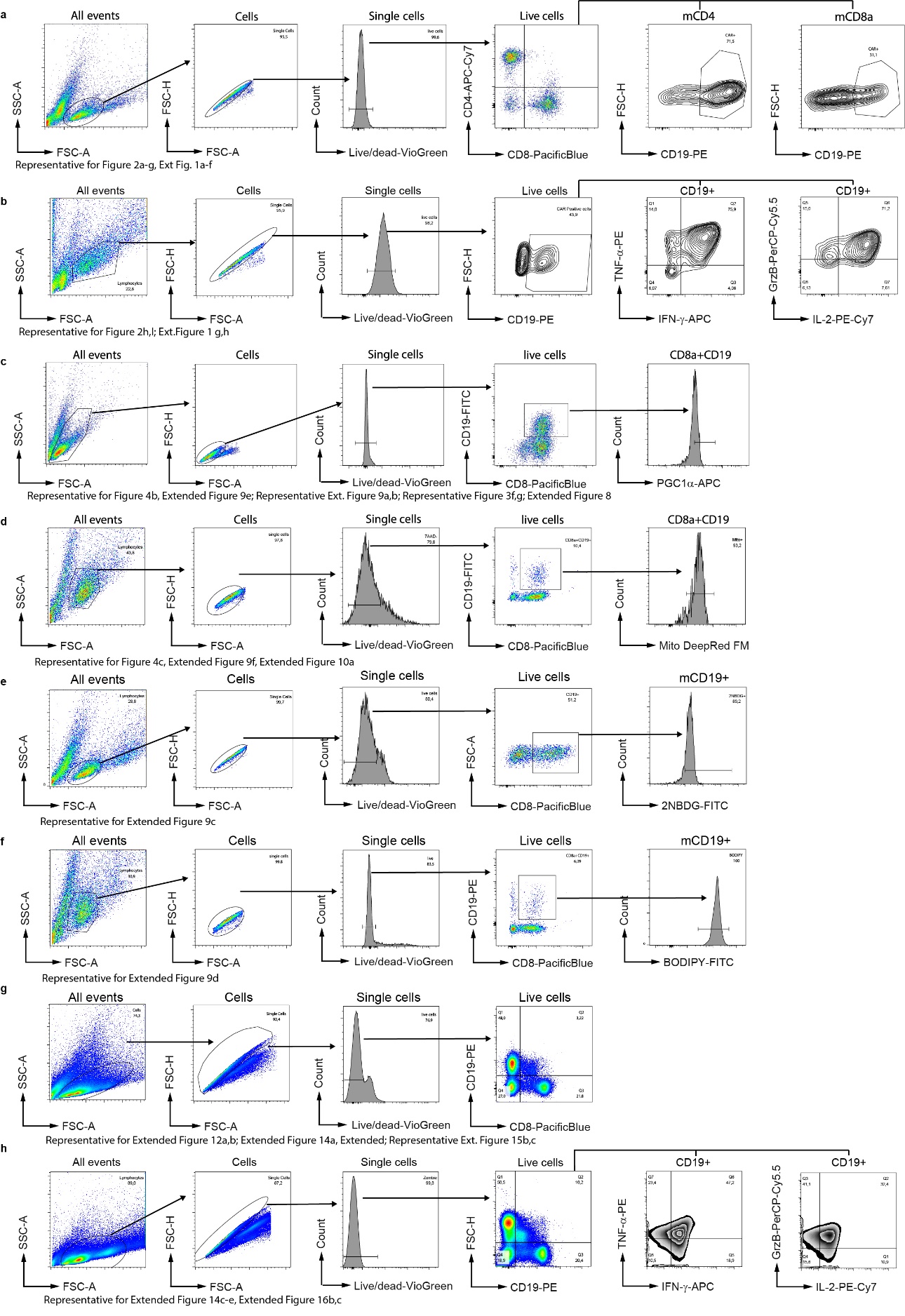

**Extended Figure 20: Gating Strategies. a** - **h**, Exemplary gating strategies for flow cytometry analysis of *in vitro* and *ex vivo* analyzed T cells.
