## Supplementary Tables for "Metabolization of microbial postbiotic pentanoate drives anti-cancer CAR T cells"

**Microbial metabolite-guided CAR T cell engineering enhances anti-tumor immunity via epigenetic-metabolic crosstalk**

Staudt et al.

**Supplemetary Tables**

**Supplementary Table 1: Patient characteristics - German cohort**

| parameter | **n (%)** |
| --- | --- |
| **total** | 66 (100) |
| **disease** |  |
| high-grade B-NHL | 43 (64) |
| multiple myeloma | 11 (16) |
| follicular lymphoma | 3 (4) |
| solid tumor | 3 (4) |
| mantle cell lymphoma | 2 (3) |
| acute lymphoblastic leukemia | 2 (3) |
| PCNSL | 2 (3) |
| **CAR-T cell product** |  |
| Axicabtagene ciloleucel | 32 (48) |
| Tisagenlecleucel | 14 (21) |
| Idecabtagene vicleucel | 11 (17) |
| experimental CAR | 5 (8) |
| Brexucabtagene autoleucel | 4 (6) |
| **CRS**, total | 61 (92) |
| - grade 1 | 25 (38) |
| - grade 2 | 30 (45) |
| - grade 3 | 5 (8) |
| - grade 4 | 1 (2) |
| **ICANS**, total | 14 (21) |
| - grade 1 | 4 (6) |
| - grade 2 | 5 (8) |
| - grade 3 | 2 (3) |
| - grade 4 | 3 (5) |
| **Any antibiotic therapy*** | 20 (30) |
| -PIM | 14 (21) |
| -Non-PIM antibiotics | 6 (9) |
| *Abbreviations:*  B-NHL = B-Non-Hodgkin lymphoma, PCNSL = primary central nervous system B cell lymphoma, CAR = chimeric antigen receptor, CRS = cytokine release syndrome, ICANS = immune effector cell-associated neurotoxicity syndrome, PIM = piperacillin/tazobactam, imipenem, meropenem. *Antibiotic therapy excluding pneumocystis prophylaxis as per institutional standards | |

**Supplementary Table 2: Patient characteristics – Memorial Sloan Kettering Cancer Center Cohort**

| Characteristic | N = 60 |
| --- | --- |
| Median Age (IQR) | 69 (57, 73) |
| Sex |  |
| Female | 21 (35%) |
| Male | 39 (65%) |
| Diagnosis |  |
| Large B-cell Lymphoma | 58 (97%) |
| Follicular Lymphoma | 2 (3.3%) |
| CAR-T product |  |
| Axicabtagene ciloleucel | 38 (63%) |
| Lisocabtagene maraleucel | 6 (10%) |
| Tisagenlecleucel | 16 (27%) |
| PIM exposure |  |
| Not exposed before infusion | 51 (85%) |
| Exposure before infusion | 9 (15%) |
| Max grade CRS |  |
| 0 | 11 (18%) |
| 1 | 19 (32%) |
| 2 | 27 (45%) |
| 3 | 3 (5.0%) |
| Max grade ICANS |  |
| 0 | 40 (67%) |
| 1 | 6 (10%) |
| 2 | 7 (12%) |
| 3 | 7 (12%) |

IQR = Interquartile Range, CAR-T = Chimeric Antigen Receptor T-cell, PIM = Piperacillin-Tazobactam, Imipenem, Meropenem, CRS = Cytokine Release Syndrome, ICANS = Immune Effector Cell-Associated Neurotoxicity Syndrome.

**Supplementary Table 3: Antibody list**

| **Reagent/Resource** | **Source** | **Identifier (Cat-number)** | **Clone** |
| --- | --- | --- | --- |
| Goat polyclonal anti-hamster IgG | MP Biomedicals | Cat#: 56984 | N/A |
| Ultra-LEAF™ Purified anti-mouse CD3ε Antibody | Biolegend | Cat#: 100359 | 145-2C11, RRID:AB_2616673 |
| Ultra-LEAF™ Purified anti-mouse CD28 Antibody | Biolegend | Cat#: 102116 | 37.51, RRID:AB_11147170 |
| Anti-mouse CD3 Pacific blue™ | Biolegend | Cat#: 100214 | 17A2, RRID:AB_493645 |
| Anti-mouse CD3 PE-Cyanine7 | Biolegend | Cat#: 100220 | 17A2, RRID:AB_1732057 |
| Anti-mouse CD4 FITC | BioLegend | Cat#: 100406 | GK1.5, RRID:AB_312691 |
| Anti-mouse CD4 APC-Cyanine7 | BioLegend | Cat#: 100414 | GK1.5, RRID:AB_312699 |
| Anti-mouse CD8a APC-Cyanine7 | BioLegend | Cat#: 100714 | 53-6.7, RRID:AB_312753 |
| Anti-mouse CD8a Pacific blue™ | BioLegend | Cat#: 100725 | 53-6.7, RRID:AB_493425 |
| Anti-mouse CD8a PE-Cyanine7 | BioLegend | Cat#: 100722 | 53-6.7, RRID:AB_312761 |
| Anti-mouse CD45 Pacific blue™ | BioLegend | Cat#: 103126 | 30-F11, RRID:AB_493535 |
| Anti-mouse IFN-γ APC | Biolegend | Cat#: 505810 | XMG1.2, RRID:AB_315404 |
| Anti-mouse IFN-γ APC-Cyanine7 | Biolegend | Cat#: 505850 | XMG1.2, RRID:AB_2616698 |
| Anti-mouse IFN-γ  Pacific blue™ | Biolegend | Cat#: 505818 | XMG1.2, RRID:AB_893526 |
| Anti-mouse TNF-α  PerCP/Cyanine5.5 | BioLegend | Cat#: 506322 | MP6-XT22, RRID:AB_961434 |
| Anti-mouse TNF-α PE | BioLegend | Cat#: 506306 | MP6-XT22, RRID:AB_315427 |
| Anti-mouse IL-2 PE-Cyanine7 | BioLegend | Cat#: 503832 | JES6-5H4, RRID:AB_2561750 |
| Anti-human/mouse Granzyme B recombinant Pacific blue™ | Biolegend | Cat#: 372218 | QA16A02, RRID:AB_2728385 |
| Anti-human/mouse Granzyme B recombinant PerCP/Cyanine5.5 | Biolegend | Cat#: 372212 | QA16A02, RRID:AB_2728379 |
| Anti-human/mouse Granzyme B recombinant APC | Biolegend | Cat#: 372204 | QA16A02, RRID:AB_2687028 |
| PGC1a Alexa Fluor^®^ 647 | Santa Cruz | Cat#: sc-518025 | D-5, RRID:AB_2890187 |
| Anti-mouse FoxP3 PE | eBioscience | Cat#: 12-5773-82 | FJK-16s, RRID:AB_465936 |
| Anti-mouse T-bet PE | Biolegend | Cat#: 644810 | 4B10, RRID:AB_2200542 |
| Anti-CD45.1 APC | eBioscience | Cat#: 17-0453-82 | A20, RRID:AB_469398 |
| Anti-mouse IL17A PE | eBioscience | Cat#: 12‐7177‐81 | eBio17B7, RRID:AB_763582 |
| Anti-human EGFR APC | Self- conjugated | N/A | Erbitux, RRID:AB_2459632 |
| Anti-mouse CD19 PE | BioLegend | Cat#: 152408 | 1D3/CD19, RRID:AB_2629817 |
| Anti-mouse CD19 FITC | BioLegend | Cat#: 152404 | 1D3/CD19, RRID:AB_2629813 |
| Anti-mouse CD19 PerCP/Cyanine5.5 | BioLegend | Cat#: 152406 | 1D3/CD19, RRID:AB_2629815 |
| 7AAD staining solution | Miltenyi | Cat#: 130-111-568 | N/A |
| Anti-mouse phospho Akt1 (Ser473) PE | Cell signaling | Cat#: 9271 | RRID:AB_329825 |
| Anti-mouse phospho STAT5 (Tyr694) APC | Invitrogen | Cat#: **17-9010-42** | SRBCZX, RRID:AB_2573272 |
| Donkey anti-rabbit IgG Alexa Fluor® 647 | Biolegend | Cat#: 406414 | Poly4064, RRID:AB_2563202 |
| Anti-Rabbit IgG Alexa Fluor™ 546 | Invitrogen | Cat#: **A-11035** | RRID:AB_143051 |
| Acetyl-Histone H3 (Lys27) | Cell signaling | Cat#: 8173 | D5E4, RRID: AB_10949503 |
| Acetyl-Histone H3 (Lys9/Lys14) | Cell signaling | Cat#: 9677 | RRID: AB_1147653 |
| Anti-mouse phospho mTOR (Ser2448) PE | Invitrogen | Cat#: **12-9718-42** | MRRBY, RRID:AB_2572724 |
| Anti-mouse phospho S6 (Ser235/236) PE | Invitrogen | Cat#: **12-9007-42** | cupk43k, RRID:AB_2572667 |
| Isotype rabbit IgG | Cell signaling | Cat#: 2729 | RRID:AB_1031062 |

### Isotype controls

| **Reagent/Resource** | **Source** | **Identifier (Cat-number)** | **Clone** |
| --- | --- | --- | --- |
| Rat IgG2a, κ APC/Cyanine 7 Isotype control | Biolegend | Cat#: 400524 | RTK2758 |
| Rat IgG2a, κ Pacific blue Isotype control | Biolegend | Cat#: 400527 | RTK2758 |
| Rat IgG2a, κ PE/Cyanine 7 Isotype control | Biolegend | Cat#: 400522 | RTK2758 |
| Rat IgG2a, κ PE Isotype control | Biolegend | Cat#: 400508 | RTK2758 |
| Rat IgG2a, κ PerCP/Cyanine5.5 Isotype control | Biolegend | Cat#: 400532 | RTK2758 |
| Rat IgG2a, κ FITC Isotype control | Biolegend | Cat#: 400506 | RTK2758 |
| Mouse IgG2a, k Alexa Flour 647 Isotype control | Santa Cruz | Cat#: sc-3878 | RRID:AB_737242 |
| Rat IgG2b, κ pacific blue™ Isotype control | Biolegend | Cat#: 400627 | RTK4530 |
| Rat IgG2b, κ; PE/Cyanine7 Isotype control | Biolegend | Cat#: 400618 | RTK4530 |
| Rat IgG2b, κ FITC Isotype control | Biolegend | Cat#: 400634 | RTK4530 |
| Rat IgG2b, κ APC/Cyanine 7 Isotype control | BioLegend | Cat#: 400624 | RTK4530 |
| Rat IgG2a, κ APC/Cyanine 7 Isotype control | Biolegend | Cat#: 400524 | RTK2758 |
| Rat IgG1, κ APC Isotype control | Biolegend | Cat#: 400412 | RTK2071 |
| Rat IgG1, κ APC/Cyanine7 Isotype control | Biolegend | Cat#: 400422 | RTK2071 |
| Rat IgG1, κ Pacific blue™ Isotype control | Biolegend | Cat#: 400419 | RTK2071 |
| Rat IgG1, κ PerCP/Cyanine5.5 Isotype control | Biolegend | Cat#: 400426 | RTK2071 |
| Rat IgG1, κ PE Isotype control | Biolegend | Cat#: 400408 | RTK2071 |
| MouseIgG1, κ APC Isotype control | Biolegend | Cat#: 400120 | MOPC-21 |
| MouseIgG1, κ Brilliant violet 421™ Isotype control | Biolegend | Cat#: 400157 | MOPC-21 |
